# Prenylation of axonally translated proteins controls NGF-dependent axon growth

**DOI:** 10.1101/2020.01.21.914606

**Authors:** Emily Scott-Solomon, Rejji Kuruvilla

**Affiliations:** Department of Biology, Johns Hopkins University, 3400 N. Charles St, 227 Mudd Hall, Baltimore, Maryland 21218, USA

**Keywords:** Compartmentalized signaling in neurons, protein lipidation, local protein synthesis, axons, neurotrophin trafficking, neuronal development

## Abstract

Compartmentalized signaling is critical for cellular organization and specificity of functional outcomes in neurons. Here, we report that post-translational lipidation of newly synthesized proteins in axonal compartments allows for short-term and autonomous responses to extrinsic cues. Using conditional mutant mice, we found that protein prenylation is essential for sympathetic axon innervation of target organs. We identify a localized requirement for prenylation in sympathetic axons to promote axonal growth in response to the neurotrophin, Nerve Growth Factor (NGF). NGF triggers prenylation of proteins including the Rac1 GTPase in axons, counter to the canonical view of prenylation as constitutive, and strikingly, in a manner dependent on axonal protein synthesis. Newly prenylated proteins localize to TrkA-harboring endosomes in axons, and promote receptor trafficking necessary for axonal growth. Thus, coupling of prenylation to local protein synthesis presents a mechanism for spatially segregated cellular functions during neuronal development.

## Introduction

Spatial partitioning of biochemical processes is a fundamental principle that underlies cellular structure and specificity of functional responses in all cells, but is particularly relevant in polarized nerve cells. Neurons rely on the asymmetric distribution of RNA, proteins, and lipids to specialized sub-cellular domains to accomplish compartment-specific functions. Proteins involved in growth cone migration, axon extension, and neurotransmitter release are enriched in axons, whereas proteins involved in post-synaptic functions including neurotransmitter receptors accumulate in dendrites and spines. How such segregation of cellular material is established and maintained in neuronal compartments to allow autonomous responses to extrinsic cues or neural activity remains poorly defined.

Lipidation is a post-translational modification that makes proteins hydrophobic and facilitates their insertion into the plasma membrane or intracellular membranes. Protein prenylation is an irreversible modification that involves the transfer of farnesyl or geranylgeranyl isoprenoid lipids to conserved carboxyl terminal CaaX motifs in proteins, and is predicted to affect at least 200 mammalian proteins (Wang and Casey, 2016). Despite critical functions of proteins predicted to be prenylated in cellular signaling, cytoskeleton remodeling, and vesicular trafficking, the functional relevance of the prenyl groups for individual proteins is poorly understood. Further, protein prenylation is considered to be a constitutive process that occurs ubiquitously throughout the cytoplasm in eukaryotic cells (Sinensky, 2000; Wang and Casey, 2016). The highest expression of isoprenoid lipids and prenyl transferases, enzymes responsible for adding isoprenoid lipids to newly synthesized proteins, is found in the nervous system (Joly et al., 1991; Tong et al., 2008). Given their complex morphologies and cellular polarity, prenylation could be particularly critical for spatially segregating protein functions in neurons.

Here, we describe a mechanism where a neurotrophic factor couples local synthesis of protein effectors with their lipid modification in axonal compartments to allow acute and spatial responses necessary for axon development. Using compartmentalized cultures of sympathetic neurons, we identified a unique need for local protein prenylation in axons to promote growth in response to Nerve Growth Factor (NGF), a target-derived axon growth and survival factor for sympathetic neurons. NGF acutely triggers prenylation of proteins in distal axons and growth cones of sympathetic neurons. Notably, the lipid modifications occur on proteins that are locally synthesized in axons. The newly modified proteins localize to endosomes harboring TrkA receptors for NGF in axons and promote receptor trafficking, which is a critical determinant of trophic signaling. In mice, protein prenylation is essential for NGF-dependent sympathetic axon innervation of targets and neuronal survival. Together, these results suggest that coupling of local protein synthesis with post-translational lipidation in axons is a mechanism for compartmentalized responses to spatial cues during neuronal development.

## Results

### Protein geranylgeranylation is required locally in axons for NGF-dependent axon growth

To determine where protein prenylation occurs in polarized neurons, we investigated the expression of farnesyl and geranylgeranyl transferase I (FTase and GGTase I, respectively) in peripheral sympathetic neurons. FTase and GGTase I catalyze the addition of either a farnesyl or geranylgeranyl isoprenoid lipid to proteins. They are expressed as heterodimers which share a common α-subunit and have distinct β-subunits. Immunostaining with an antibody against the shared prenyl transferase α-subunit (Luo et al., 2003) showed robust expression in sympathetic neuronal cell bodies and axon fibers innervating a target field, the salivary glands, in mice at postnatal day 5 **(Figures S1A, B).** This is a developmental period when sympathetic neurons rely on the neurotrophin, NGF, released from peripheral targets, for their survival and axon innervation (Glebova and Ginty, 2005). Immunostaining of cultured sympathetic neurons revealed prenyl transferase expression in cell bodies, axons, and growth cones (**Figure S1C).** Immunoblotting of lysates from compartmentalized neuron cultures, where a Teflon-grease diffusion barrier separates cell bodies from axons, showed a protein of the predicted size (44 kDa) in both compartments (**Figure S1D).** The specificity of the antibody was verified in PC12 cells by shRNA transfection and immunoblotting (**Figure S1E**). In sympathetic neuron cell bodies and PC12 cells, two higher molecular weight bands at approximately 60-65 kDa were also observed that were reduced by shRNA-mediated knockdown, suggesting a likely post-translational modification. Together, these results indicate that the molecular machinery required for protein prenylation is present in cell bodies, axons, and even growth cones of sympathetic neurons.

To visualize protein prenylation in sympathetic neurons, we developed a live-cell feeding assay in compartmentalized cultures. NGF (50 ng/ml) was added only to distal axonal compartments, recapitulating the release of neurotrophins from target tissues. Prenylation was visualized by incubating either cell bodies or axons with a membrane-permeable prenyl lipid analog, propargyl-farnesol (isoprenoid analog), which is metabolically incorporated into cellular proteins at native CaaX sites using endogenous prenyl transferase activity (DeGraw et al., 2010). Newly modified proteins were visualized by conjugation to a fluorophore (biotin azide-streptavidin-Alexa-488) using click chemistry-based labeling in fixed cells. We observed prominent isoprenoid reporter labeling, indicative of newly prenylated proteins, appearing in a punctate or sometimes tubular pattern, in cell bodies, axon shafts, and distal axons **(Figure 1A-C).** The non-diffuse labeling pattern, specifically in axons, suggests that protein prenylation does not occur throughout the cytoplasm, but rather in discrete sub-cellular sites.

**Figure 1.**
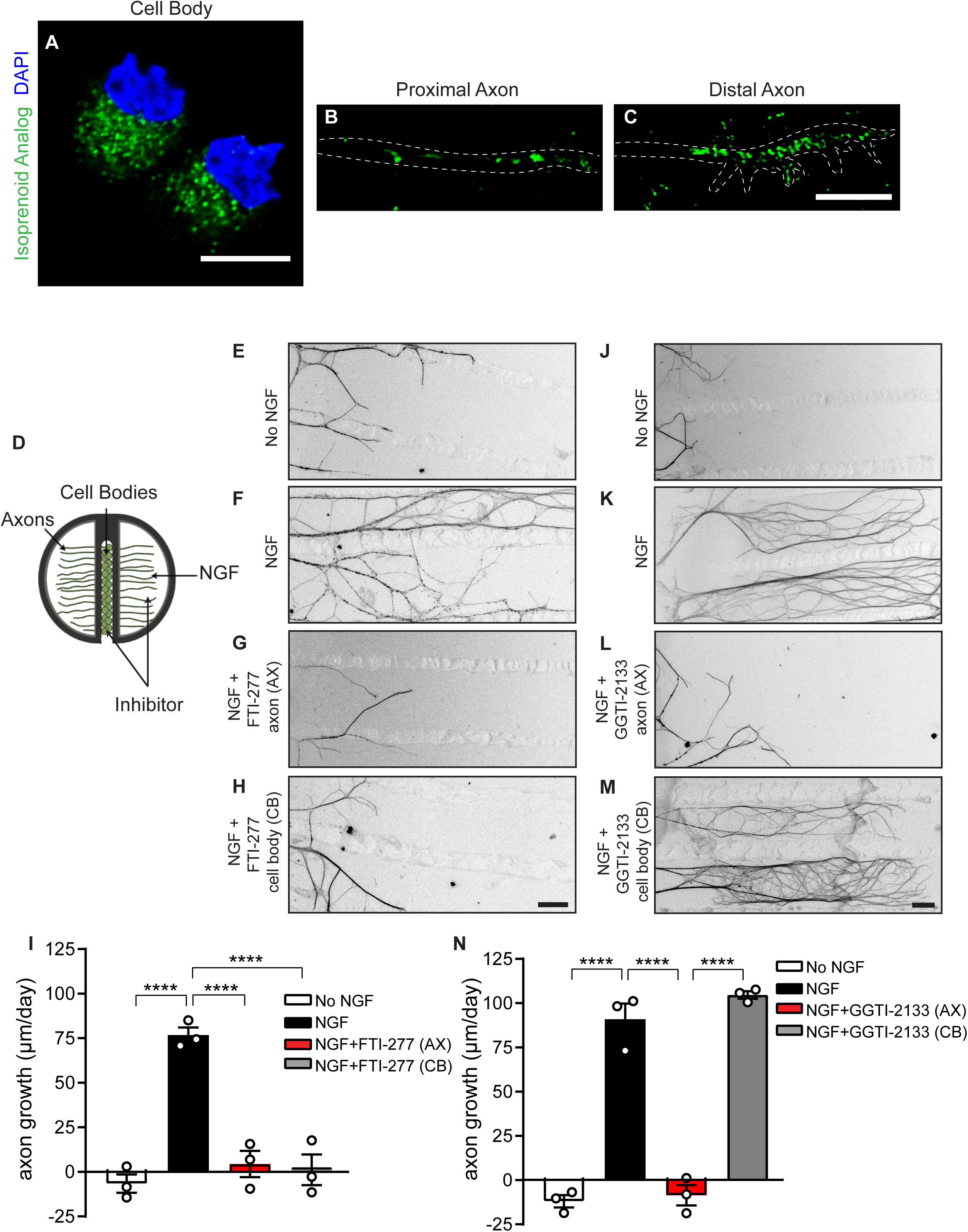
Protein geranylgeranylation is required locally in axons for NGF-dependent axon growth. **(A-C)** Visualization of newly prenylated proteins in cell bodies **(A)**, proximal axons **(B)**, and distal axons **(C)**, of sympathetic neurons by live-feeding with a membrane-permeant isoprenoid analog and click labeling. Sympathetic neurons were grown in compartmentalized chambers with NGF (50 ng/ml) added to distal axon compartments. **(D)** In compartmentalized cultures of sympathetic neurons, cell bodies and distal axons are in fluidically isolated compartments, allowing spatially-restricted inhibition of protein prenylation. **(E-H)** Farnesylation is required in both cell bodies (CB) and axons (AX) to mediate NGF-dependent growth. Axon growth is blocked by treating axons **(G)** or cell bodies **(H)** with the FTase inhibitor (FTI-2177, 100 nM). **(I)** Average growth rate of axons measured at 24-hr intervals for 72 hr following FTI-2177 treatment. Data presented as mean ± s.e.m from n=3 experiments with at least 20 axons traced per condition. ****p<0.0001, one-way ANOVA Tukey’s multiple comparisons. **(J-M)** NGF-dependent axon growth is abolished by addition of the GGTase I inhibitor (GGTI-2133, 75 nM) to axon **(L)**, but not, cell body **(M)** compartments. **(N)** Quantification of axon growth with GGTI-2133 treatment. Results are mean ± s.e.m (n=3 experiments). ****p<0.0001 one-way ANOVA Tukey’s multiple comparisons. NGF (50 ng/ml) was added only to distal axon compartments. Panels are representative images of axons immunostained with anti-β-III-tubulin 72 hr after adding inhibitors. Scale bars 10 μm **(A-C),** 100 μm **(E-H and J-M)**.

NGF mediates growth of sympathetic axons by binding to TrkA receptors in axons and promoting the formation of NGF-TrkA signaling endosomes that signal locally or are retrogradely trafficked to cell bodies (Scott-Solomon and Kuruvilla, 2018). Several protein effectors involved in NGF-mediated signaling, trafficking, and cytoskeletal remodeling are predicted to be prenylated (Delcroix et al., 2003; Wang and Casey, 2016; Wu et al., 2007; Zweifel et al., 2005), although the functional requirement for the lipid modifications is unknown. To address whether local protein prenylation is necessary for mediating axon growth, cell bodies or axons of neurons grown in compartmentalized cultures were treated with the membrane-permeable competitive inhibitors against FTase (FTI-277) or GGTase I (GGTI-2133) in conjunction with NGF treatment to distal axons **(Figure 1D)**. FTI-277 and GGTI-2133 are peptidomimetics that mimic CaaX motifs specific for FTase or GGTase-I respectively (Clapp et al., 2013; Lerner et al., 1995). NGF stimulation of distal axons resulted in robust growth of sympathetic axons (**Figures 1E, F, J, K, I, and N)**. The FTase inhibitor added to either cell bodies or axons attenuated NGF-dependent axon growth **(Figures 1G, H, I)**. Treatment of axons with the GGTase I inhibitor also diminished axon growth in response to NGF **(Figures 1L, N)**. Surprisingly, inhibition of GGTase I activity in cell bodies had no effect **(Figures 1M, N)**. These results indicate a specific need for proteins to undergo geranylgeranylation locally in axons to mediate NGF-dependent axon growth.

### NGF acutely triggers protein prenylation in axons

The axon-specific requirement for GGTase I in NGF-directed axon growth raised the possibility it may be regulated by NGF signaling. A fluorescent assay to measure GGTase I enzymatic activity was performed after exposing distal axons of compartmentalized sympathetic neuron cultures to NGF for 30 minutes (**Figure 2A**). NGF treatment triggered a 4-fold increase in GGTase I activity in sympathetic axons, but had no effect on enzyme activity in cell bodies within this time period **(Figure 2B)**. To visualize NGF-induced prenylation, we fed the isoprenoid analog to sympathetic neurons in the presence or absence of NGF for 4 hr. Excess analog was washed from neurons and newly modified proteins visualized by conjugation of a biotin tag and streptavidin Alexa-488 labeling. NGF increased prenylation of proteins locally in axons, particularly in axonal growth cones, compared to unstimulated neurons **(Figures 2C-G)**. No background fluorescence was observed when the isoprenoid analog was omitted in the click reaction **(Figures 2D, G)**, confirming the specificity of the prenylation signal in **Figure 2E**. Strikingly, treatment of neurons with GGTI-2133 abolished incorporation of isoprenoid analog in NGF-treated neurons **(Figures 2F, G)**, suggesting that the proteins modified by NGF treatment are GGTase I substrates. Together, these results indicate that NGF acutely regulates GGTase I activity and protein geranylgeranylation in both axons and growth cones of sympathetic neurons.

**Figure 2.**
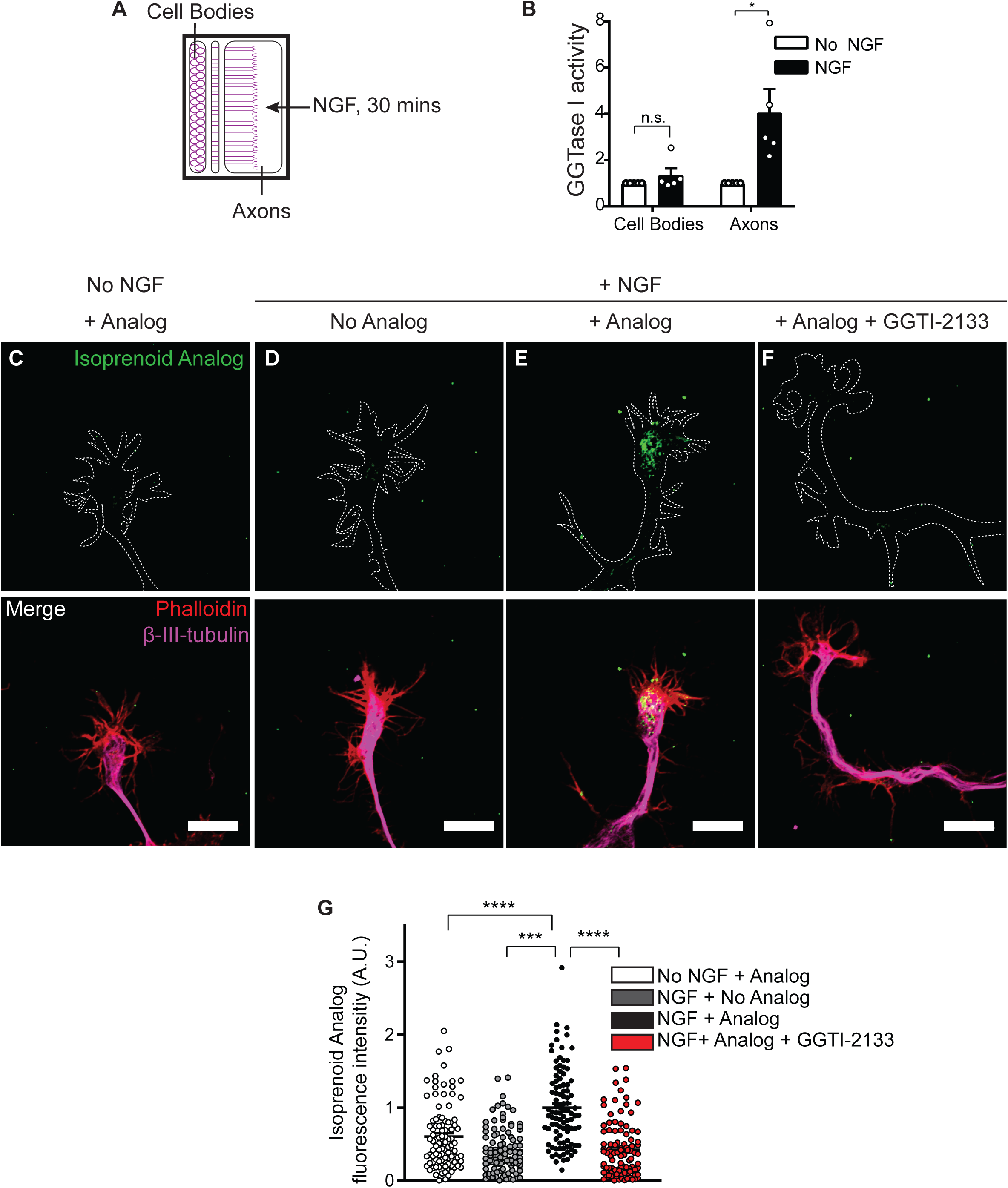
NGF acutely promotes geranylgeranylation of proteins in axons. **(A)** Compartmentalized sympathetic neurons were stimulated with NGF (50 ng/ml) added to distal axons for 30 minutes. Cell body and axon lysates were subjected to a fluorescence-based assay to measure GGTase I enzymatic activity. **(B)** NGF acutely stimulates GGTase I activity in sympathetic axons. GGTase I activity was normalized to total protein amount, n=5 independent experiments, *p<0.05, n.s. not significant, t-test. **(C-F)** NGF induces geranylgeranylation of proteins in growth cones of sympathetic axons. Enhanced incorporation of the isoprenoid analog in NGF-treated neurons (50 ng/ml, 4 hr) compared to unstimulated neurons, which is abolished by addition of GGTI-2133 (75 nM). Newly prenylated proteins were visualized by live-labeling with isoprenoid analog, followed by conjugation to biotin-streptavidin-488. No background fluorescence is observed when the isoprenoid analog is excluded from the click reaction in panel C. Growth cones were visualized by phalloidin labeling and anti-β-III-tubulin immunostaining. Scale bar: 10 μm. (**G**) Quantification of isoprenoid fluorescence intensity normalized to F-actin signal in growth cones. Data represent mean ± s.e.m from n=5 experiments with at least 20 growth cones analyzed per condition, ***p<0.001, ****p<0.0001, one-way ANOVA Tukey’s multiple comparisons.

### Protein prenylation is required for TrkA trafficking in axons

NGF signaling is initiated by binding TrkA receptors in sympathetic axons, receptor dimerization and autophosphorylation, and internalization of ligand/receptor complexes in endosomes where internalized TrkA receptors continue to signal (Scott-Solomon and Kuruvilla, 2018). TrkA endosomes acutely regulate signaling pathways in axons, and are also retrogradely transported to cell bodies to activate transcriptional programs needed for long-term axonal growth and neuron survival (Scott-Solomon and Kuruvilla, 2018). TrkA trafficking is known to rely on several effector proteins, especially small GTPases, that are predicted to be prenylated (Harrington et al., 2011; Kawata et al., 1990; Wu et al., 2001). Based on the appearance of newly prenylated proteins in discrete punctae or tubules **(Figures 1B, C)**, we asked whether the modified proteins localize to endosomal compartments in axons, and in particular, to TrkA-harboring endosomes. To monitor trafficking of surface TrkA receptors in sympathetic neurons, we utilized a chimeric Trk receptor-based live-cell antibody feeding assay (Ascano et al., 2009). Neurons were infected with an adenoviral vector expressing FLAG-tagged chimeric Trk receptors that have the extracellular domain of TrkB, and the transmembrane and intracellular domain of TrkA (Ascano et al., 2009). Sympathetic neurons do not normally express TrkB and chimeric receptors respond to the TrkB ligand, Brain-Derived Neurotrophic Factor (BDNF), but retain the signaling properties of TrkA. Surface chimeric FLAG-TrkB:A receptors were live labeled with anti-FLAG antibody. Neurons were stimulated with BDNF (50 ng/ml, 4 hr) in the presence of the isoprenoid analog, following which the remaining surface-bound anti-FLAG antibody stripped with mild acid washes. Ligand stimulation increased the intracellular accumulation of chimeric Trk receptors (**Figures 3A-C, top panel, and 3F**) and protein prenylation in axons (**Figures 3A-C, second panel from top**). Notably, 55% of Trk receptors that underwent ligand-induced endocytosis co-localized with newly prenylated proteins in axons (**Figures 3C, third panel and 3E**). These results suggest that the proteins prenylated in response to neurotrophin stimulation accumulate, in part, on TrkA-containing endosomes in axons. GGTI-2133 treatment attenuated ligand-induced uptake of isoprenoid analog **(Figure 3D),** as expected. Surprisingly, GGTI-2133 treatment significantly diminished neurotrophin-induced intracellular accumulation of Trk receptors in sympathetic axons **(Figures 3D, top panel, and 3F).** Together, these results suggest that newly prenylated proteins are associated with TrkA-harboring endosomes in axons, and that protein prenylation likely promotes NGF-dependent axonal growth by influencing receptor trafficking.

**Figure 3.**
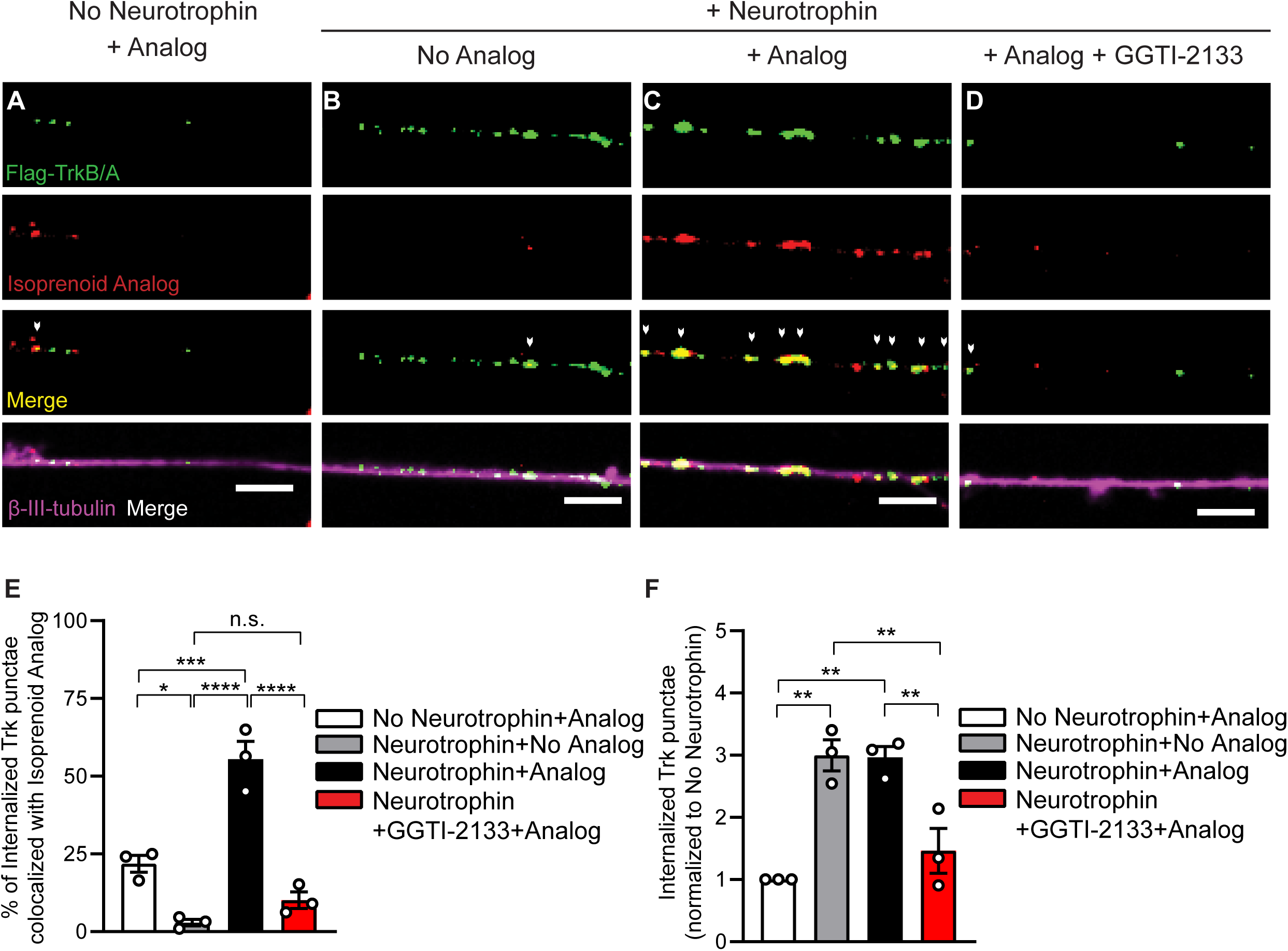
Prenylated proteins associate with Trk endosomes and promote receptor trafficking. **(A-D)** Co-localization of newly prenylated proteins with internalized Trk receptors in neurotrophin-stimulated sympathetic axons. Cultured sympathetic neurons, expressing FLAG-TrkB:A chimeric receptors, were live-labeled with FLAG antibody and isoprenyl analog under non-permeabilizing conditions and stimulated with BDNF (50 ng/ml) in the presence or absence of GGTI-2133 (75 nM) for 4 hr. Surface FLAG antibody was stripped, leaving only internalized receptors labeled. White arrows indicate co-localization between FLAG-Trk punctae and geranylgeranylation signal. Axons were visualized by β-III-tubulin immunostaining. Scale bar: 5 μm. **(E)** Percentage of internalized FLAG-Trk receptors that co-localize with prenylated proteins in axons. Data presented as mean ± s.e.m. from n=3 experiments with 15-20 axons analyzed per condition per experiment. *p<0.05, ***p<0.001, ****p<0.0001, n.s. not significant, one-way ANOVA Tukey’s multiple comparisons. **(F)** GGTase I activity is required for intracellular accumulation of surface Trk receptors in axons. Internalized receptors were calculated as number of FLAG:Trk punctae per μm. Measurements were taken from axonal segments 75-100 μm from the distal tip. Results are mean ± s.e.m. from n=3 experiments and expressed relative to the “no neurotrophin” condition. 15-20 axons were analyzed per condition per experiment. **p<0.01, one-way ANOVA Tukey’s multiple comparisons.

### NGF-induced prenylation relies on local translation in axons

The ability of NGF to promote geranylgeranylation of proteins in axons raises a question as to the source of these nascent proteins. Proteins are often prenylated immediately following synthesis (Wang and Casey, 2016). Further, our results that local protein geranylgeranylation is essential for axon growth **(Figures 1J-N)** suggests that anterograde transport of already modified proteins from cell bodies is not sufficient. Previously, screens of axonal mRNAs have identified many transcripts that are known to encode for prenylated proteins (Gumy et al., 2011). Together, these findings raised the intriguing possibility that the proteins being geranylgeranylated in axons in response to NGF are locally translated. To determine the contribution of intra-axonal protein synthesis to local geranylgeranylation in response to NGF, we used a sympathetic ganglia explant culture system that permits the mechanical removal of cell bodies leaving the axons in isolation (**Figure 4A**) (Andreassi et al., 2010). Isolated axons remain morphologically intact and respond to NGF for at least 8 hr following removal of cell bodies (Andreassi et al., 2010). After removal of cell bodies, isolated axons were incubated with the isoprenoid analog for 6 hr in the presence or absence of NGF. To determine the contribution of axonal protein synthesis to NGF-induced prenylation, incorporation of the isoprenoid analog in NGF-treated axons was assessed in the presence or absence of the translation inhibitor, cycloheximide (CHX). As seen previously, NGF promoted robust prenylation of proteins in isolated axons, which was abolished by CHX treatment **(Figures 4B-F)**. These results suggest that NGF-induced lipid modifications occur on proteins that are locally synthesized in axons.

**Figure 4.**
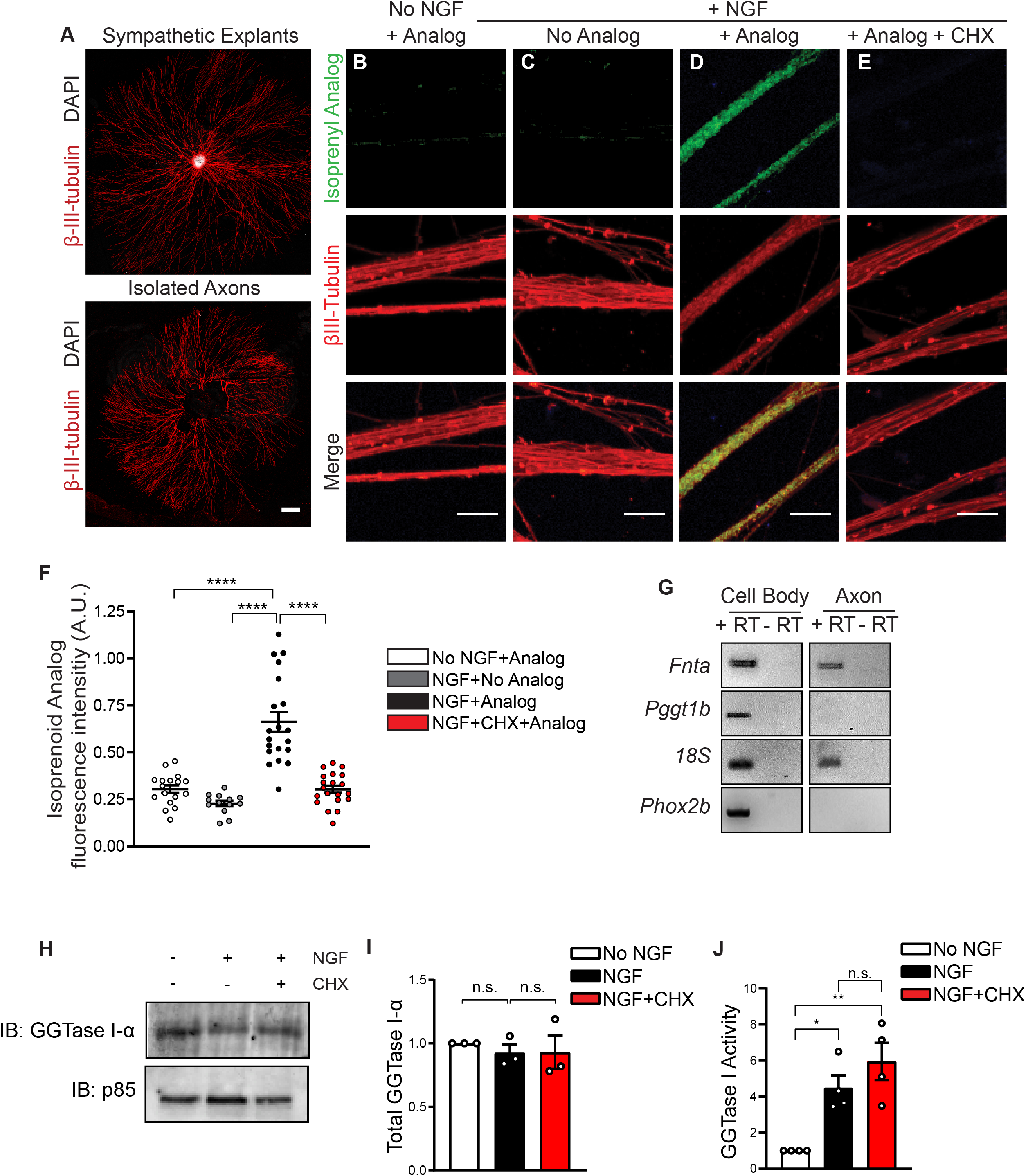
NGF-induced prenylation requires local protein synthesis in axons. **(A)** Sympathetic ganglia explant cultures where removal of cell bodies by mechanical excision allows axonal proteins to be labeled by the isoprenoid analog. β-III-tubulin immunostaining was used to visualize axons and DAPI indicates nuclei. Scale bar: 500 μm. **(B-E)** NGF (50 ng/ml) induces protein prenylation in isolated axons that is blocked by the translational inhibitor, cycloheximide (CHX 25 μM, 6 hr). Newly prenylated proteins were visualized by labeling with isoprenoid analog and conjugation to biotin-streptavidin-488. Axons were immunostained with β-III-tubulin. Scale bar: 500 μm. **(F)** Quantification of protein prenylation in isolated axons normalized to β-III-tubulin immunofluorescence. Results are mean ± s.e.m from n=10 experiments with at least 13 explants analyzed per condition, ****p<0.0001, one-way ANOVA Tukey’s multiple comparisons. **(G)** RT-PCR analysis shows presence of *Fnta* mRNA encoding for GGTase I-α, but not *Pggt1b* mRNA for GGTase I-β, in axons of compartmentalized sympathetic neurons. *18S* rRNA has been reported to be present in axons (Briese et al., 2016) and *Phox2b* is a cell body-specific transcript. (**H**) Immunoblotting shows that GGTase I-α protein levels in axons are unaffected by NGF (50 ng/ml) or NGF+CHX (25 μM) treatment over 6 hr. Immunoblotting for the p85 subunit of PI-3K was used for normalization of protein amounts. **(I)** Densitometric quantification of GGTase I-α levels normalized to p85. Results are mean ± s.e.m from n=3 experiments, 50-70 explants pooled per condition for each experiment. (**J**) NGF-induced increase in GGTase I enzymatic activity is independent of protein synthesis. Sympathetic neurons were stimulated with NGF (50 ng/ml) in the presence of CHX (25 μM) for 6 hr. Protein lysate was subjected to a fluorescence-based assay to measure GGTase I enzymatic activity. GGTase I activity was normalized to total protein amounts. Results are mean ± s.e.m. from n=4 experiments and expressed relative to the “no neurotrophin” condition. *p<0.05, **p<0.01, n.s. not significant, one-way ANOVA Tukey’s multiple comparisons.

One explanation for the dependence of axonal prenylation on local protein synthesis could be that the prenylation enzymes themselves are locally translated in axons. To test this prediction, we performed RT-PCR analysis of mRNA isolated from distal axons of sympathetic neurons grown in compartmentalized cultures, and observed that transcripts encoding for the GGTase I-α subunit (*Fnta*), but not the GGTase I-β sub-unit (*Pggt1b*), was localized in sympathetic axons (**Figure 4G**). However, GGTase I-α protein levels were unaffected by NGF treatment or NGF+CHX treatments for 6 hr in isolated sympathetic axons (**Figure 4H-I**). To directly determine if NGF-induced GGTase I enzymatic activity relied on protein synthesis, we assessed the effect of CHX on GGTase-I activity in NGF-treated sympathetic neurons. NGF stimulation enhanced GGTase I enzymatic activity as observed previously **(Figure 2B)**, and this activation was not affected by CHX treatment (**Figure 4J**). Together, these results suggest that the reliance of axonal prenylation on local translation is due to the synthesis of GGTase I substrates, and not GGTase I itself.

### NGF induces prenylation of Rac1 in axons

Which proteins are locally lipid-modified in sympathetic axons in response to NGF? The Rac1 GTPase is a known GGTase I substrate (Roberts et al., 2008). Rac1 is also a key effector of NGF trophic signaling, TrkA trafficking and axonal morphology (Harrington et al., 2011) making it an attractive candidate for NGF-regulated prenylation in axons. To assess local Rac1 geranylgeranylation, isolated sympathetic axons were incubated with the isoprenoid analog in the presence or absence of NGF for 6 hr. Rac1 was immunoprecipitated from axon lysates, conjugated to a TAMRA azide tag using click chemistry, and prenylated Rac1 was detected by immunoblotting with an anti-TAMRA antibody. We observed a robust 2-fold increase in prenylated Rac1 in axons in response to NGF stimulation (**Figures 5A, B**). Treatment of axons with the GGTase I inhibitor abolished NGF-induced Rac1 prenylation (**Figures 5A, B**) without affecting total Rac1 levels (**Figure 5C**). Thus, NGF induces axonal geranylgeranylation of Rac1. We next asked if prenylation of Rac1 in axons was dependent on local synthesis. RT-PCR analysis of mRNA isolated from distal axons of sympathetic neurons grown in compartmentalized cultures demonstrated that *Rac1* mRNA was detected in axons (**Figure S2A**). To determine if Rac1 geranylgeranylation was dependent on local synthesis, isolated axons were incubated with the isoprenoid analog and stimulated with NGF in the presence or absence of CHX. Prenylated Rac1 was detected by click labeling and immunoblotting. Treatment of isolated axons with CHX suppressed the NGF-induced increase in Rac1 prenylation **(Figures 5D, E)**, and also reduced axonal levels of Rac1 protein (**Figure S2B**). Together, these results indicate that NGF-induced prenylation of Rac1 in axons is dependent on local translation. Of note, NGF treatment enhanced Rac1 prenylation by 2-2.5-fold even after accounting for increased axonal Rac1 levels elicited by NGF (**Figures 5B and 5E)**. These data support that NGF-induced increase in Rac1 prenylation is not merely due to enhanced Rac1 protein synthesis, but also by regulation of GGTase I enzymatic activity (see **Figures 2B and 4J)**. Thus, NGF regulates the local synthesis of Rac1 in axons <u>and</u> also ensures its post-translational modification to enable axonal functions.

**Figure 5.**
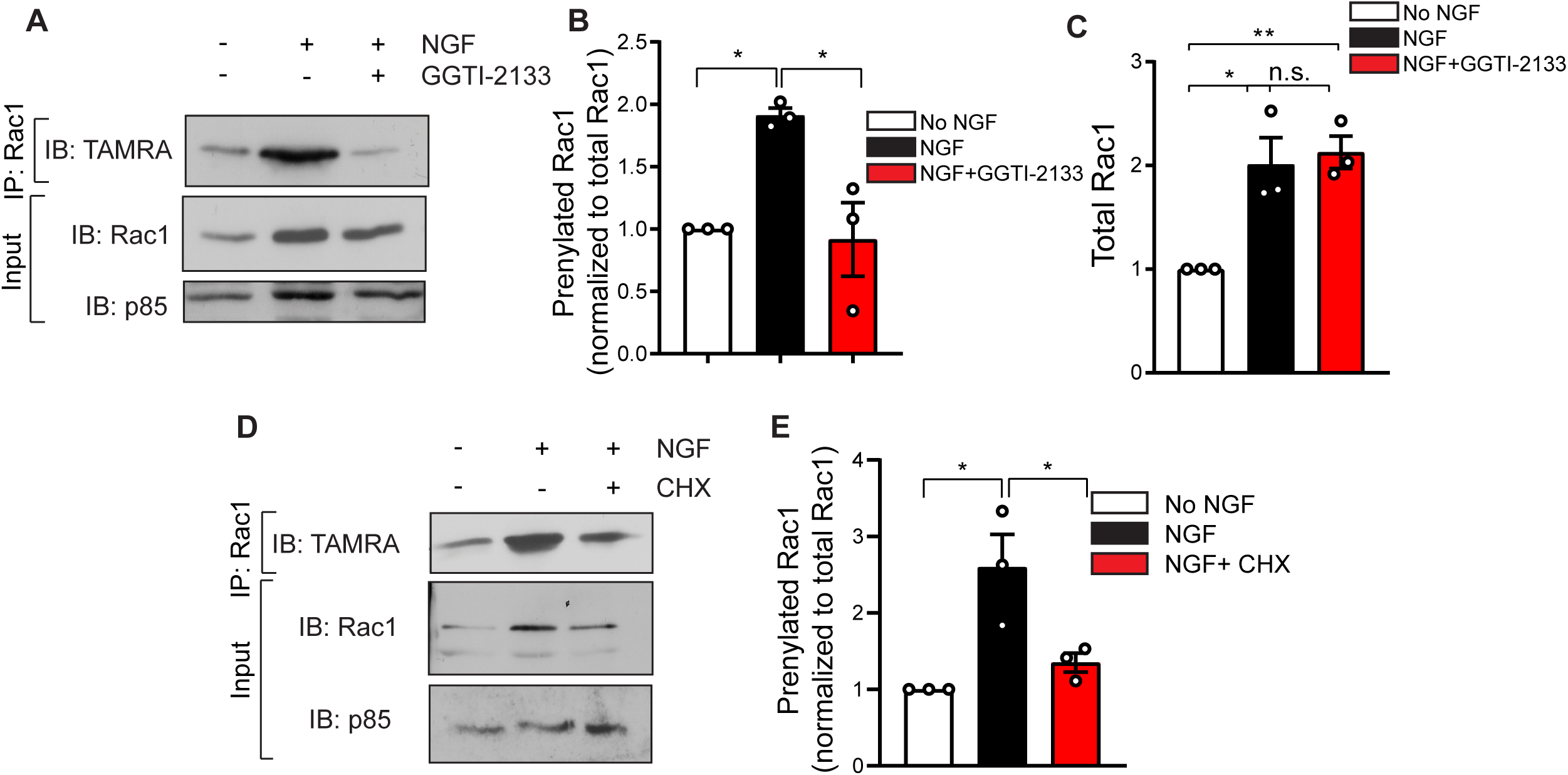
NGF induces prenylation of axonally translated Rac1. **(A)** Rac1 is geranylgeranylated in axons in response to NGF. Isolated sympathetic axons were labeled with the isoprenoid analog in the presence of NGF, NGF+GGTI-2133, or anti-NGF (no NGF) for 6 hr. Prenylated Rac1 was detected by Rac1 immunoprecipitation, conjugation to a TAMRA azide tag, and immunoblotting with an anti-TAMRA antibody. Supernatants were immunoblotted for Rac1, followed by stripping and re-probing for p85 to normalize for protein amounts. **(B)** Quantification of prenylated Rac1 normalized to total Rac1 protein in axons. Results are mean ± s.e.m from n=3 experiments, 50-70 explants pooled per condition for each experiment, *p<0.05, n.s. not significant, one-way ANOVA Tukey’s multiple comparisons. **(C)** NGF-induced increase in axonal Rac1 protein is not affected by GGTase I inhibition. Quantification of total Rac1 in axons normalized to p85. Results are mean ± s.e.m from n=3 experiments, 50-70 explants pooled per condition for each experiment, *p<0.05, **p<0.01, one-way ANOVA Tukey’s multiple comparisons. **(D)** NGF-induced Rac1 prenylation depends on axonal protein synthesis. Isolated axons were labeled with isoprenoid analog in the presence of anti-NGF, NGF, or NGF+CHX for 6 hr. Newly prenylated Rac1 was detected by immunoprecipitation, conjugation to a TAMRA-azide tag, and TAMRA immunoblotting. Supernatants were immunoblotted for Rac1. Rac1 immunoblots were stripped and re-probed for p85 to control for loading. **(E)** NGF-induced increase in prenylation is attenuated by CHX treatment. Quantification of prenylated Rac1 normalized to total Rac1 protein in axons. Results are mean ± s.e.m. from n=3 experiments, 50-70 explants pooled per condition for each experiment, *p<0.05, n.s. not significant, one-way ANOVA Tukey’s multiple comparisons.

### Prenylation of axonally translated Rac1 promotes NGF-dependent axon growth

Asymmetric localization of mRNA transcripts depends on cis-elements commonly located within their 3’UTRs (Andreassi et al., 2018). To study the intra-axonal synthesis of Rac1 and to identify element(s) responsible for *Rac1* mRNA trafficking to axons, we performed rapid amplification of 3’cDNA ends (3’RACE) on mRNA isolated from either cell bodies or distal axons of sympathetic neurons grown in compartmentalized cultures. Interestingly, 3’RACE analysis revealed two isoforms of *Rac1* mRNA with different 3’UTR lengths; a long 3’UTR (∼1500 bp) and a short 3’UTR (∼250 bp) (**Figures 6A, B and Figure S3A**). Both isoforms have the same coding sequence. The sequence of the short isoform aligned to the first 250 bp of the long isoform upstream of a poly-adenylation site (**Figure 6B)**, suggesting that the two isoforms likely arise from alternative polyadenylation. Although both *Rac1* isoforms were found in cell bodies of sympathetic neurons, only the *Rac1* long 3’UTR isoform was detected in axons **(Figure 6A and Figure S3A)**. This finding for *Rac1* is consistent with previous reports for other mRNA isoforms where a long 3’UTR biases for localization to neuronal processes(Andreassi et al., 2018). To visualize the sub-cellular distribution of protein products of the two *Rac1* isoforms with distinct 3’UTRs, we generated constructs containing mCherry-tagged coding sequence of *Rac1* fused to either long (mCherry-Rac1-L) or short (mCherry-Rac1-S) *Rac1* 3’UTR. Sympathetic neurons were electroporated either with mCherry-Rac1-L or mCherry-Rac1-S, and after 36 hr, the mCherry fluorescence visualized and quantified in cell bodies, proximal, and distal axons. While both constructs were expressed at similar levels in cell bodies and proximal axons (**Figures 6C-E**), the mCherry-Rac1-L protein was selectively enriched (3-fold increase) in distal axons (**Figures 6C-D, F**). These results suggest that axonal localization of *Rac1* mRNA is accomplished via the long 3’UTR, and imply that Rac1-L is the major contributor to the prenylated protein in sympathetic axons.

**Figure 6.**
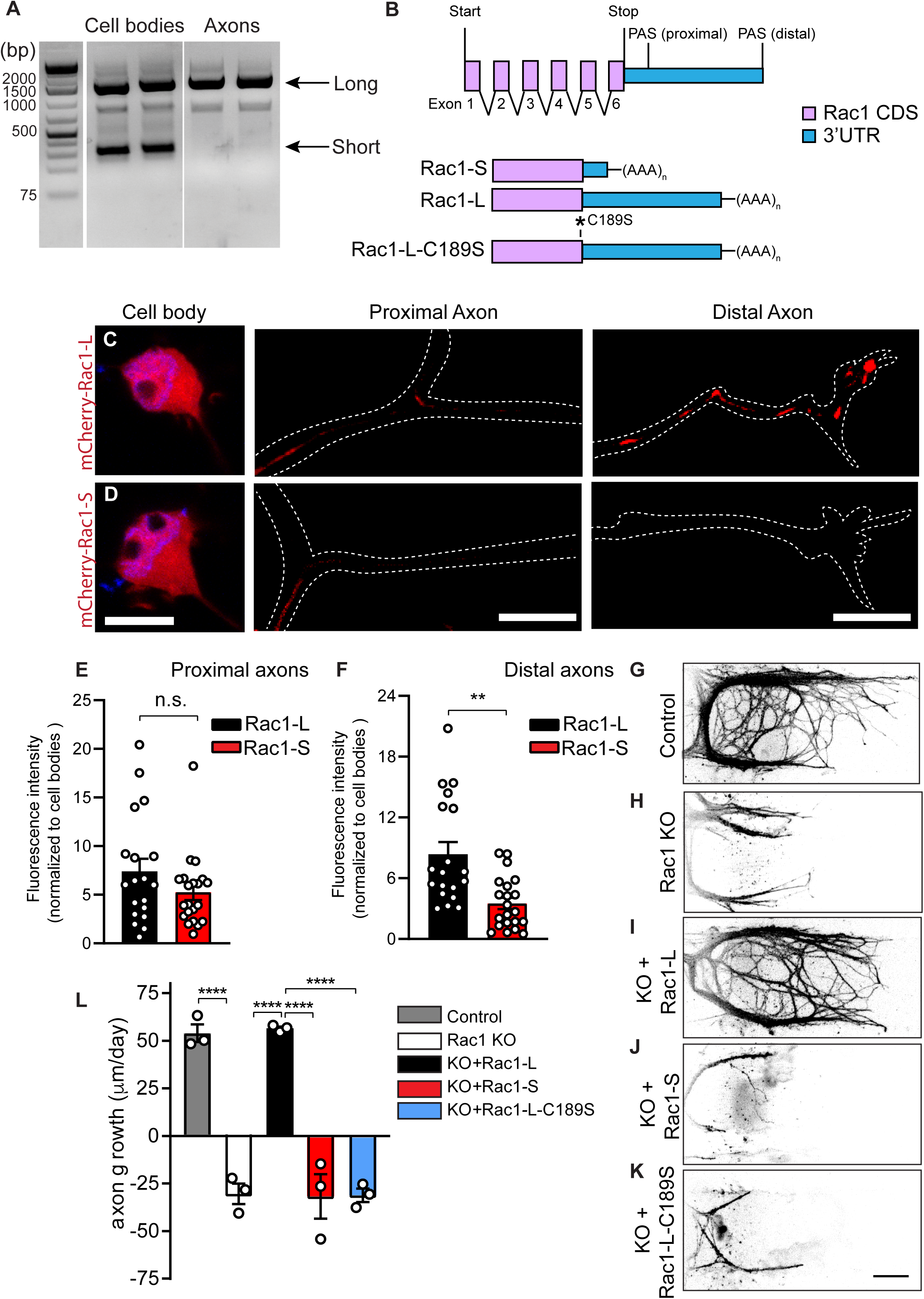
Prenylation of axonally translated Rac1 directs NGF-dependent axon growth. **(A)** 3’RACE analysis on mRNA isolated from compartmentalized cultures of sympathetic neurons reveals two *Rac1* isoforms with distinct 3’UTR lengths; an isoform with a short 3’UTR that is specific to cell bodies, and an isoform with a long 3’UTR found in both cell bodies and axons. **(B)** *Rac1* 3’UTR has two unique polyadenylation sites (PAS), suggesting that the two isoforms likely arise from alternative polyadenylation. Schematic representing adenoviral constructs expressing Rac1-long 3’UTR (Rac1-L), Rac1-short 3’UTR (Rac1-S), or Rac1-long 3’UTR with the CaaX motif mutated (Rac1-L-C189S). **(C-D)** Axonal localization of *Rac1* mRNA is mediated by the long 3’UTR. Sympathetic neurons were electroporated with plasmids containing mCherry-tagged coding sequence of human *Rac1* fused to either the long or short 3’UTR of rat *Rac1*. Neurons were cultured for 36 hr before imaging. m-Cherry-Rac1-L is found in cell bodies, proximal and distal axons, while mCherry-Rac1-S is restricted to cell bodies and proximal axons. Axons are outlined in images with white dashed lines. Scale bar: 10 μm. (**E**) Similar levels of mCherry-Rac1 fluorescence were observed in proximal axons of neurons expressing Rac1-L and Rac1-S. **(F)** mCherry-Rac1 fluorescence was significantly enriched in distal axons of neurons expressing Rac1-L compared to Rac1-S. mCherry fluorescence was measured for a distance of 75-100 um for proximal axons (0 um position is at soma) or distal axons (0 um position is at the axon terminal). mCherry-Rac1 fluorescence in proximal or distal axons was normalized to soma fluorescence levels. Results are mean ± s.e.m. from n=3 independent experiments (19-21 neurons analyzed), *p<0.01, n.s. not significant, t-test. **(G-L)** NGF-mediated axon growth requires both local Rac1 synthesis and its prenylation in axons. *Rac1^fl/fl^* sympathetic neurons were cultured in compartmentalized chambers in the presence of NGF (50 ng/ml) on distal axons for 7 days to allow axonal growth into lateral compartments. Neurons were then infected with adenoviruses expressing GFP, Cre, Cre + Rac1-L, Cre + Rac1-S, or Cre + Rac1-L-C189S and grown in culture media in the absence of NGF for 36 hr. NGF was added back to distal axons, and axon growth assessed in 24 hr intervals for 48 hr. Representative images of axons immunostained with anti-β-III-tubulin 48 hr after infection. Scale bar: 100μm. **(L)** Average rate of axon extension (μm/day) assessed at 24-hr intervals for a total of 48 hr. Data presented as mean ± s.e.m. from n=3 experiments., ****p<0.0001, one-way ANOVA Tukey’s multiple comparisons.

We next asked if Rac1-S or Rac1-L was sufficient to support NGF-mediated axon growth in Rac1 knockout neurons. Sympathetic neurons isolated from *Rac1^fl/fl^* mice (Glogauer et al., 2003) at postnatal day 0.5 (P0.5) were grown in compartmentalized cultures in the presence of NGF added to distal axons for 7 days, and then infected with either an adenoviral vector expressing Cre recombinase to acutely delete *Rac1*, or LacZ as a control. Quantitative RT-PCR showed that Cre delivery elicited a significant depletion (∼82%) of *Rac1* transcript within 36 hr (**Figure S3B**). As expected, inducible deletion of Rac1 elicited axonal retraction despite the presence of NGF (**Figures 6G, H, and L**). Remarkably, expression of a Rac1-L, but not Rac1-S, adenoviral construct completely restored axon growth in response to NGF in Rac1-deleted neurons (**Figures 6I, J, and L**). This result suggests that axonal localization of *Rac1* mRNA, accomplished via the long 3’UTR, is sufficient to promote axon growth in response to NGF. Next, we asked if local prenylation of the Rac1 protein is necessary for axon growth. To prevent modification of the Rac1 isoform present in axons, the cysteine residue in the CaaX prenylation motif in the Rac1-L isoform was mutated to a serine (Rac1-L-C189S) **(Figure 6B)**. Mutation of the Rac1 prenylation motif did not affect its ability to be expressed, as determined by western blotting of mass cultures of sympathetic neurons infected with Rac1-L-C189S adenoviral vector (**Figure S3C**). Notably, analysis of NGF-dependent axon growth in compartmentalized neurons revealed that expression of Rac1-L-C189S was unable to rescue axon growth deficits in Rac1 knockout neurons (**Figures 6K and L**), in contrast to Rac1-L that had an intact prenylation motif (**Figures 6I and L**). Together, these results indicate that intra-axonal synthesis and geranylgeranylation of Rac1 is required for its function in promoting NGF-dependent axon growth.

### Protein geranylgeranylation is essential for target innervation in mice

We next addressed the relevance of protein geranylgeranylation for NGF-dependent development of the sympathetic nervous system *in vivo*. We asked if conditional deletion of *Pggt1b*, the gene coding for GGTase-I, would compromise NGF-dependent axon innervation of peripheral target tissues. To generate mice lacking GGTase I in sympathetic neurons, *Pggt1b^fl/fl^* mice (Sjogren et al., 2007) were crossed with transgenic mice expressing CRE recombinase under the control of the Dopamine Beta-Hydroxylase promoter (Parlato et al., 2007), which resulted in efficient deletion of *Pggt1b* (**Figure S4A**). Sympathetic axon innervation of peripheral organs was analyzed in newborn mice (P0.5) using whole mount tyrosine hydroxylase (TH) immunostaining in iDISCO-cleared tissues followed by light-sheet microscopy. We observed decreased sympathetic axon innervation of the kidney and heart in homozygous *DBH-Cre;Pggt1b^fl/fl^* mice and in heterozygous *DBH-Cre;Pggt1b^fl/+^* mice with one functional copy of *Pggtt1b*, compared to control litter-mates (**Figures 7A-F**). Quantification of sympathetic innervation revealed that both total axonal length and branching were affected (**Figures 7G-J**). Given that NGF also supports sympathetic neuron survival, we assessed neuronal numbers in the superior cervical ganglia (SCG), the largest and most-rostral ganglia in the sympathetic chain, in control and GGTase I mutant mice at P0.5. We observed significant cell loss in the homozygous mutants (**Figures 7K, M, and N)**, whereas heterozygous mice had normal numbers of sympathetic neurons (**Figures 7K, L, and N)**. Together, these results indicate that GGTase I is essential for the development of sympathetic neurons at the time when they are most dependent on NGF for axon growth and survival. NGF is secreted by the target tissues and acts locally in axons to support growth (Scott-Solomon and Kuruvilla, 2018). The NGF-TrkA signal is also retrogradely transported to the cell bodies, where it promotes the transcription of genes required for cell survival and axon growth (Scott-Solomon and Kuruvilla, 2018). Our findings that GGTase I is haplo-insufficient for axon innervation, but not for neuronal survival, suggest that lowered GGTase I levels primarily affect axon growth *in vivo*.

**Figure 7.**
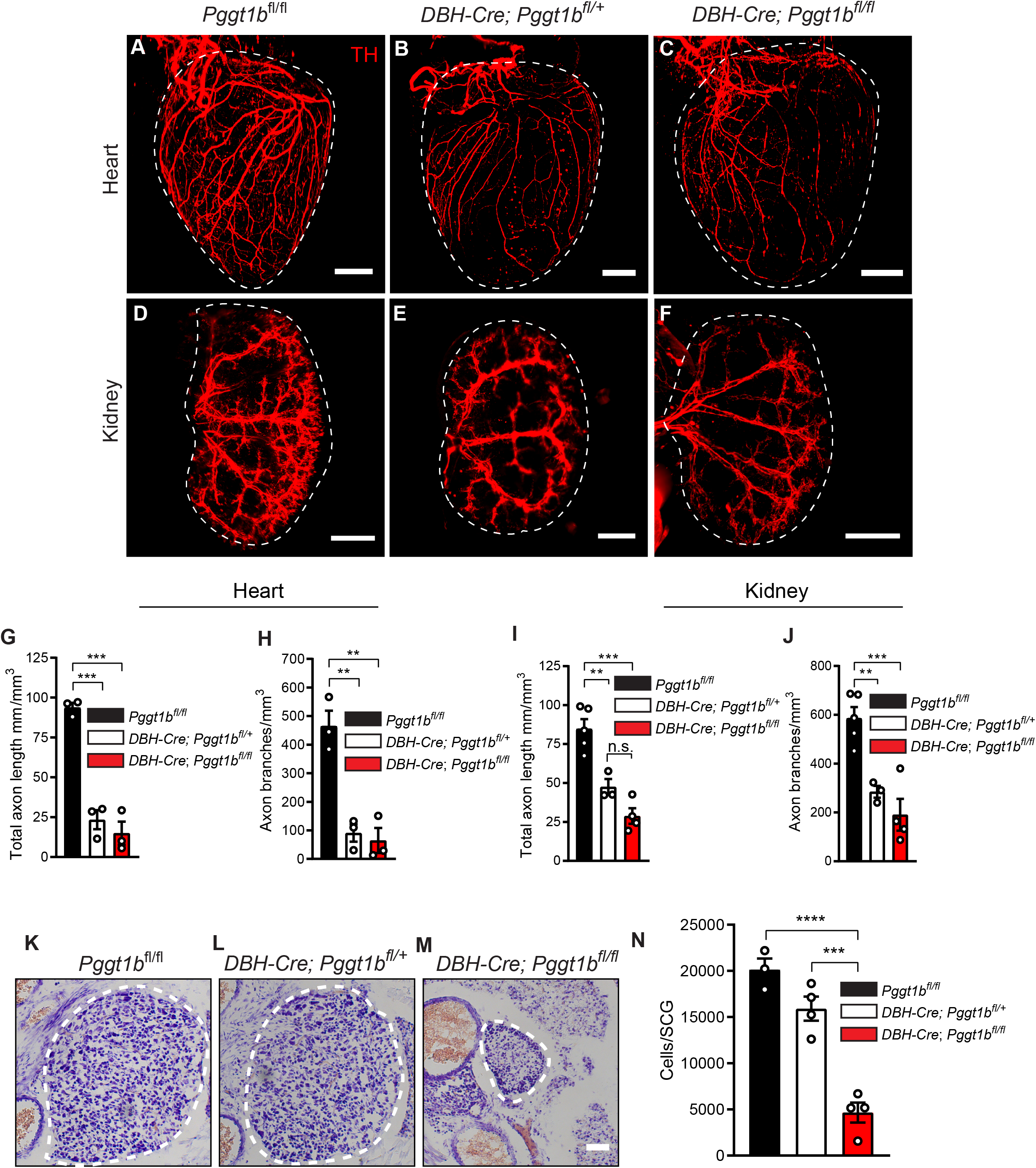
GGTase I is essential for sympathetic axon innervation of target organs *in vivo*. **(A-J)** GGTase I is essential for sympathetic axon innervation of target organs *in vivo*. iDISCO-based tissue clearing and wholemount tyrosine hydroxylase (TH) immunostaining of organs show reduced sympathetic innervation of the heart **(A-C)** and kidney **(D-F)** in heterozygous *DBH-Cre:Pggt1b^fl/+^* and homozygous *DBH-Cre:Pggt1b^fl/f^*^l^ P0.5 mice, compared to litter-mate *Pggt1b^fl/fl^* controls. **(G-J)** Quantification of axon length and branches in the heart (**G, H)** and kidney **(I,J)**. Data presented as mean ± s.e.m. from n=5 control, 3 heterozygous, and 4 homozygous mice. **p<0.01, ***p<0.001, n.s. not significant, one-way ANOVA Tukey’s multiple comparisons. **(K-N)** Sympathetic neuron numbers in the superior cervical ganglia (SCG) are reduced in homozygous, but not heterozygous, GGTase I conditional knockout mice. Counts of cell number were performed on Nissl stained tissue sections from P0.5 *DBH-Cre;Pggt1b^fl/fl^*, *DBH-Cre;Pggt1b^fl/+^* mice, and *Pggt1b^fl/fl^* littermates. Scale bar: 50 μm. Data are presented as mean ± s.e.m. from n=4 heterozygous, n=4 homozygous, and n=3 control mice. ***p<0.001, ****p<0.0001, one-way ANOVA Tukey’s multiple comparisons.

## Discussion

Asymmetrical mRNA localization and local protein synthesis is a conserved mechanism for spatio-temporal regulation of gene expression in all eukaryotic cells, but particularly relevant in neurons given their complex morphologies and extreme polarity. In neurons, specific mRNAs encoding for cytoskeleton-regulatory proteins, cellular signaling, metabolic enzymes, ion channels, synaptic proteins, and membrane biogenesis are transported to axons or dendrites where they can be rapidly translated in response to extrinsic signals such as neurotrophic factors, guidance cues, synaptic activity, and following injury (Jung et al., 2014). However, how these newly synthesized proteins are then post-translationally modified, regulated, and anchored to allow compartmentalized functions has been poorly defined. Here, we describe a mechanism where a neurotrophic factor couples local translation of protein effectors with their lipid modification in axonal compartments to allow acute and spatial responses necessary for axon development (**Figure 8**). We found that protein prenylation (geranylgeranylation) is essential for axons of sympathetic neurons to innervate their peripheral organs during development in mice. Surprisingly, geranylgeranylation is required in a compartment-specific manner in axons to promote growth in response to the neurotrophin, NGF. We show that NGF acutely triggers protein prenylation in axons and even growth cones, counter to the canonical view of prenylation as being constitutive. Remarkably, we discovered that NGF-induced lipid modifications occur on proteins that are newly synthesized in axons. We identify the Rac1 GTPase as a protein that is locally synthesized and prenylated in axons to promote NGF-dependent axonal growth. Together, these results suggest that axonal translation and geranylgeranylation might be a mechanism to enrich and regulate the local actions of the newly synthesized proteins by directing membrane association, trafficking, protein-protein interactions, and stability.

**Figure 8.**
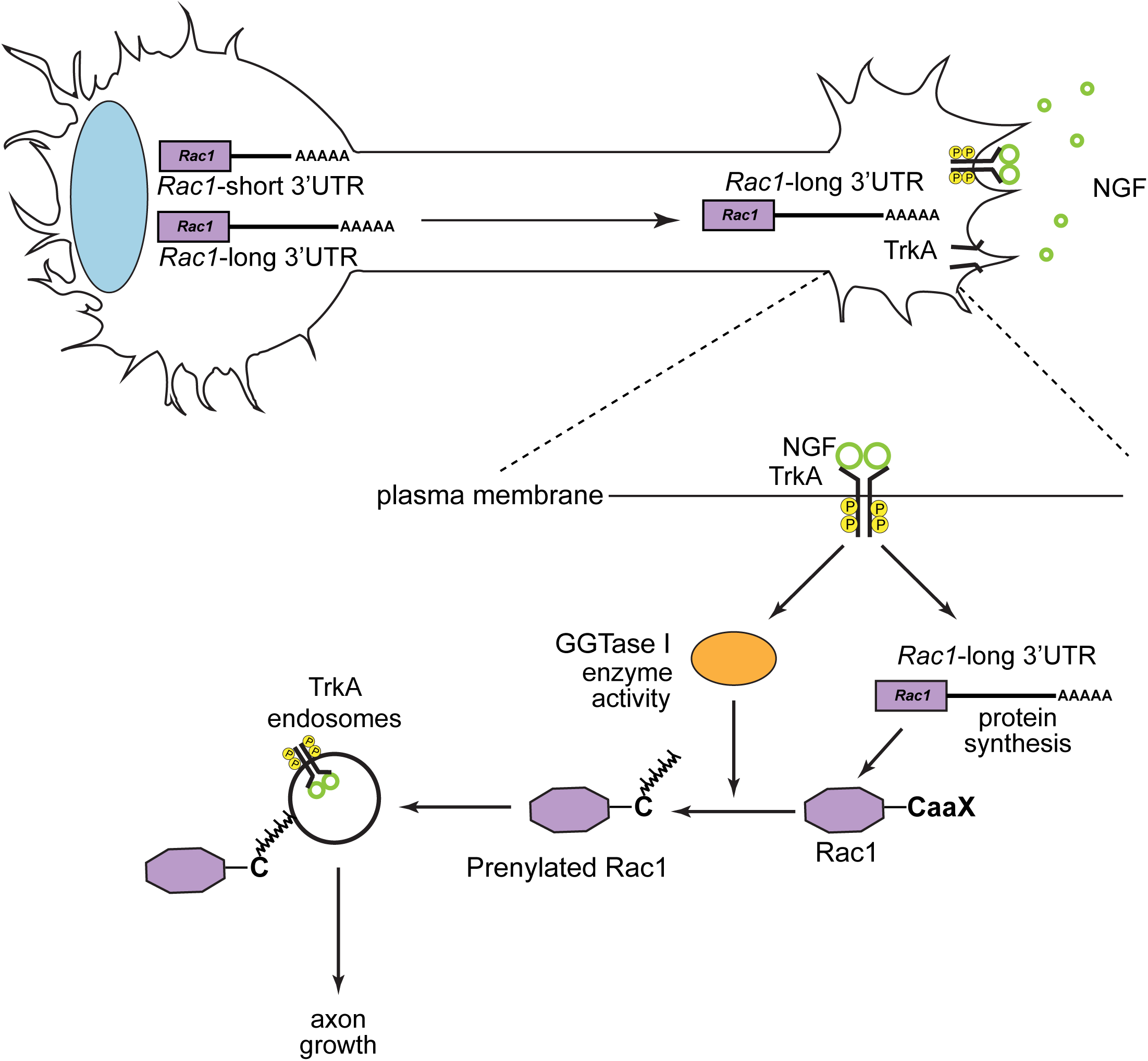
NGF couples local protein synthesis to geranylgeranylation in axons to promote axonal growth. mRNAs for growth-promoting proteins such as *Rac1* are transported to sympathetic axons where they are capable of being rapidly translated in response to extrinsic neurotrophic cues. Two isoforms of *Rac1* mRNA with distinct 3’UTR lengths are found in sympathetic neurons, with the transcript bearing the long 3’UTR preferentially targeted to axons. NGF-TrkA signaling stimulates Rac1 synthesis and geranylgeranylation in sympathetic axons to promote axonal growth. Prenylated proteins, including Rac1, likely mediate NGF-dependent growth by associating with TrkA signaling endosomes and influencing receptor trafficking in axons. The coupling of local translation to post-translational lipidation in axons is a mechanism to localize and enrich protein effectors to ensure axon-autonomous responses to extrinsic cues.

How might NGF-induced prenylation in axons support axonal growth? One mechanism might involve regulation of endosomal trafficking of TrkA receptors, a critical determinant of trophic support of sympathetic neurons by target-derived NGF. NGF-dependent development requires internalization of NGF-TrkA complexes into signaling endosomes in axon terminals to locally promote neurite elongation and branching events (Bodmer et al., 2011; Harrington et al., 2011). We found that newly prenylated proteins accumulate on TrkA endosomes upon neurotrophin stimulation. Importantly, inhibition of geranylgeranylation attenuated intracellular accumulation of Trk receptors in sympathetic axons. This could reflect a failure of endocytosis of surface Trk receptors, or enhanced recycling or degradation of internalized receptors. Recent studies highlight a role for the Rac1 GTPase in promoting trafficking of TrkA receptors and initiating NGF trophic signaling in distal axons of sympathetic neurons (Harrington et al., 2011); Rac1 is recruited to TrkA endosomes where it helps to sever a dense cytoskeletal meshwork in axon terminals to promote TrkA trafficking. NGF-induced Rac1 prenylation might be a means for its localization to TrkA endosomes. In addition to Rac1, other prenylated proteins might impinge on TrkA trafficking and signaling in axons. Two candidate proteins are the small GTPases, Rap1 and RhoB. Rap1 undergoes geranyl-geranylation in non-neuronal cells (Kawata et al., 1990), and in NGF-responsive sensory neurons, localizes to TrkA endosomes and promotes sustained signaling from internalized receptors (Wu et al., 2001). Further, geranyl-geranylation of RhoB diverts internalized Epidermal Growth Factor Receptors (EGFRs) to a recycling pathway instead of undergoing lysosomal degradation (Wherlock et al., 2004). Assessing the broad role of prenyl-modifications of Rap1, RhoB, or other small GTPases, and their local synthesis in the internalization, post-endocytic trafficking, and axonal transport of TrkA receptors and other cargo will be of interest in future studies.

We found that protein geranylgeranylation in cell bodies of sympathetic neurons is not sufficient to promote NGF-mediated axon growth. It is possible that anterograde transport of already-modified proteins from cell bodies cannot keep pace with the rapid and continuous need for cellular material in developing axons. Alternatively, geranylgeranylation may result in retention of proteins in the cellular compartments where they are made. In support of the latter, we found that the Rac1 protein product from a soma-specific transcript with a short 3’UTR is restricted to cell bodies and proximal axons of sympathetic neurons, compared to the axonal Rac1 synthesized from mRNA with a long 3’UTR that is selectively enriched at distal tips (**Figures 6C-F**). In addition to our findings for *Rac1*, several mRNAs in neurons including *BDNF* (An et al., 2008), *Importin β1* (Ben-Tov Perry et al., 2012), *mTOR* (Terenzio et al., 2018), *Impa1* (Andreassi et al., 2010) and *Shank* (Bockers et al., 2004) have isoforms with the same coding sequence but varied 3’UTR lengths because of alternative polyadenylation signals within the 3’UTR. Notably, mRNA isoforms with the longer 3’UTRs are selectively localized in axons or dendritic processes by localization elements within their long 3’UTRs (Andreassi et al., 2018). Our results in this study suggest that alternative use of 3’UTRs of transcripts and prenylation of the locally synthesized proteins in neuronal sub-compartments could be a mechanism for spatially segregated protein functions.

The relevance of prenylation to human health has been evident in diseases such as cancer and progeria, and prenylation inhibitors are being developed for clinical use (Wang and Casey, 2016). In the nervous system, limited studies implicate aberrant protein prenylation in neurological disorders. GGTase-I levels are elevated in motor neurons derived post-mortem from human patients with early onset amyotrophic lateral sclerosis (ALS) (Li et al., 2016). Further, levels of isoprenoid lipids are increased in brains of individuals with Alzheimer’s Disease (AD) (Eckert et al., 2009), and prenyl transferase inhibitors are being considered for therapeutic use in neurodegenerative disorders (Hottman and Li, 2014). These studies, although limited, highlight the potential significance of protein prenylation in establishing and maintaining connections in the nervous system, and imply that aberrant localization, activity, or stability of prenylated proteins could contribute to neuropathology. Identification of proteins that are translated and prenylated in neuronal sub-domains will define the local prenylated proteomes that allow responsiveness to extrinsic cues or neuronal activity, and their contribution to neuronal development, maintenance, and repair following injury or disease.

## Acknowledgements

We thank H. Zhao, A. Riccio, and S. Hattar for comments on the manuscript. We thank M. Bergö (Karolinska Institutet, Sweden) for providing *Pggt1b^fl/fl^* mice, W. Tourtellotte (Cedars Sinai, USA) for *DBH-Cre* mice, and C. Gerfen (NIH, USA) for *TH-Cre* mice. We thank C-C. Chen and D. Prosser for sharing plasmids for N1 Venus pA H1 backbone and human myc-tagged Rac1 CDS, respectively. This work was supported by NIH R01 awards, NS114478 and NS107342 to R.K., and NIH Training grant T32GM007231 to E.S-S.

## Author Contributions

E.S-S. and R.K. designed the study and wrote the manuscript. E.S-S. performed the experiments and analyzed the data. R.K. contributed to data analyses.

## STAR METHODS

### KEY RESOURCES TABLE

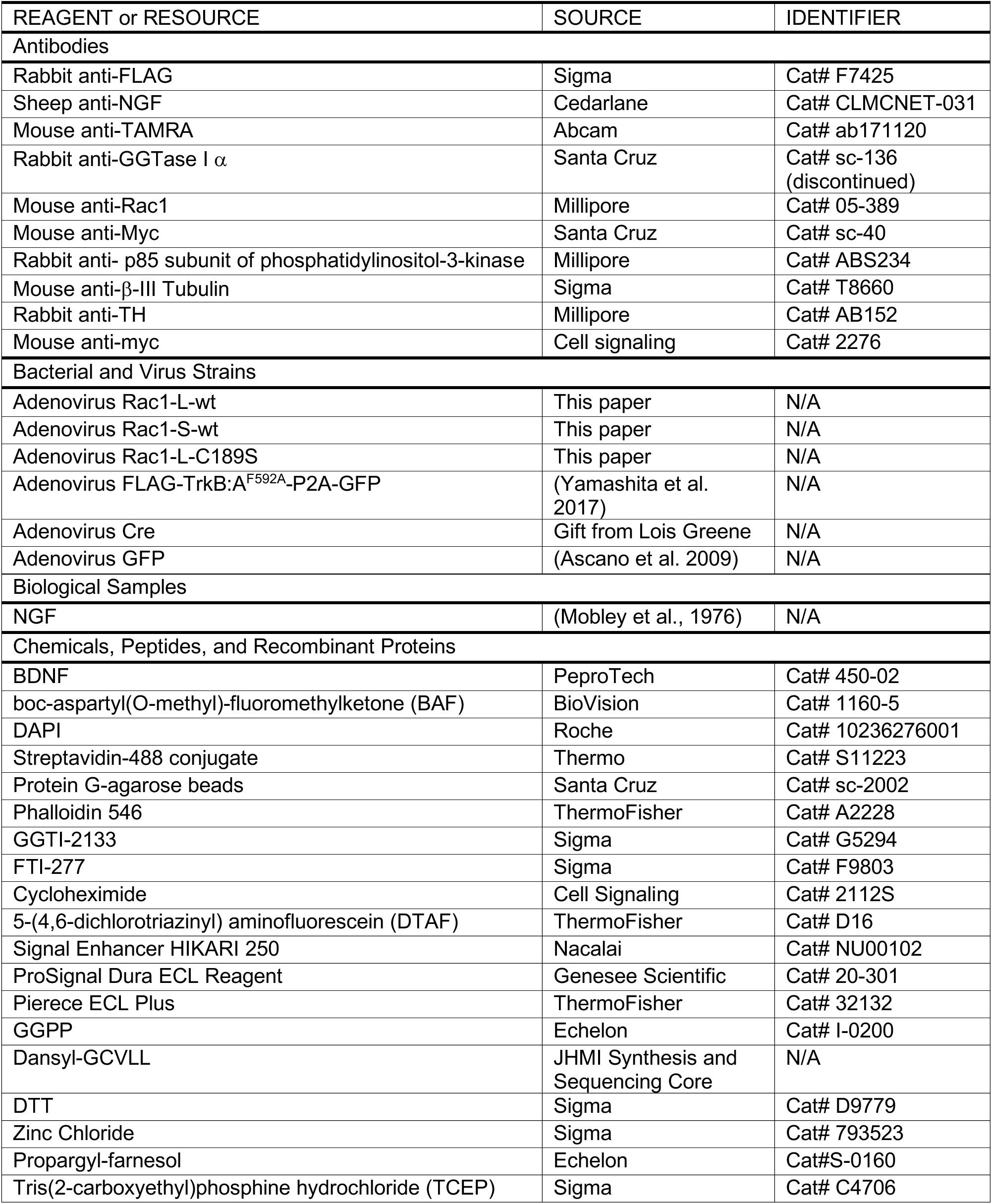
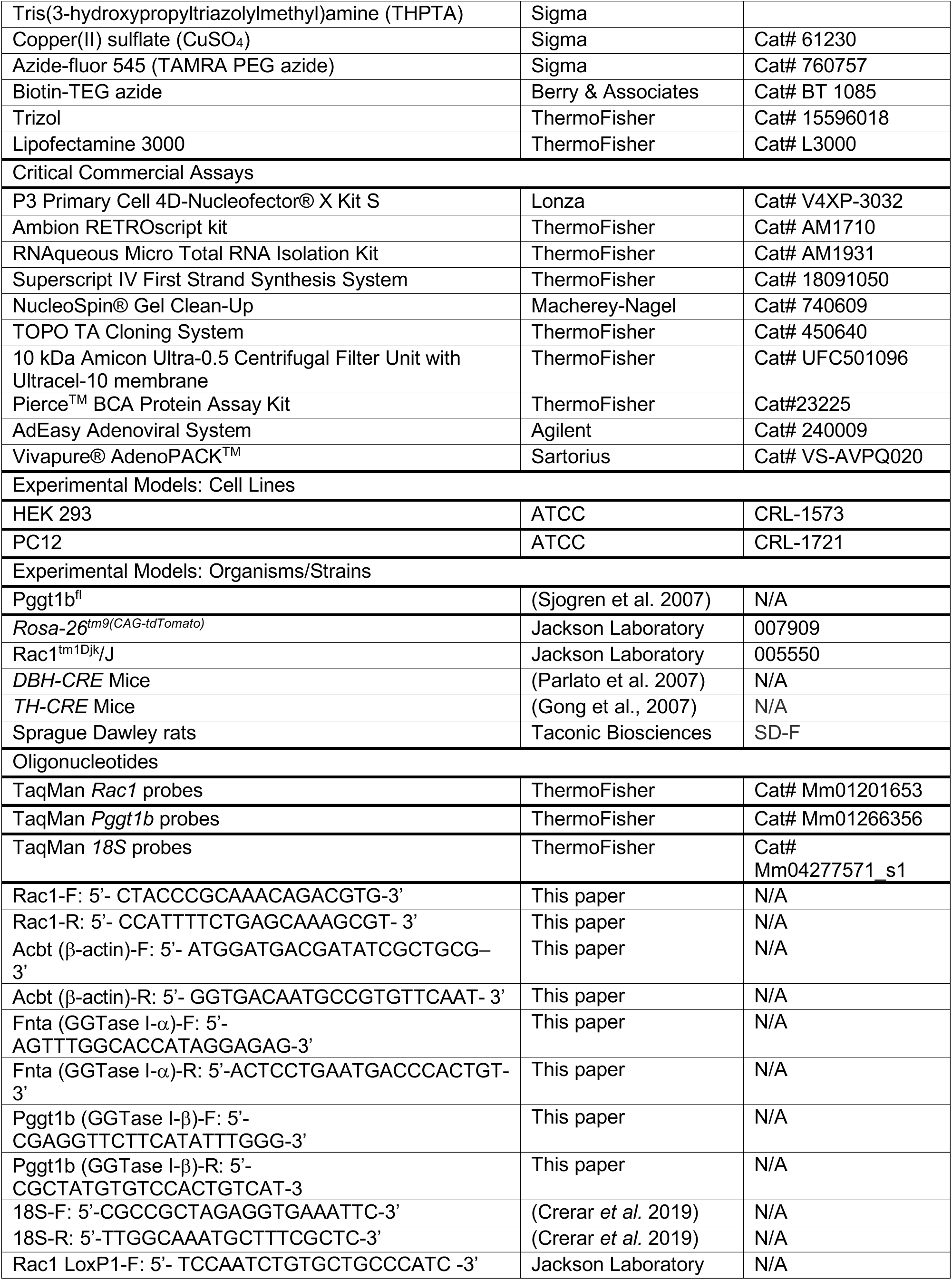
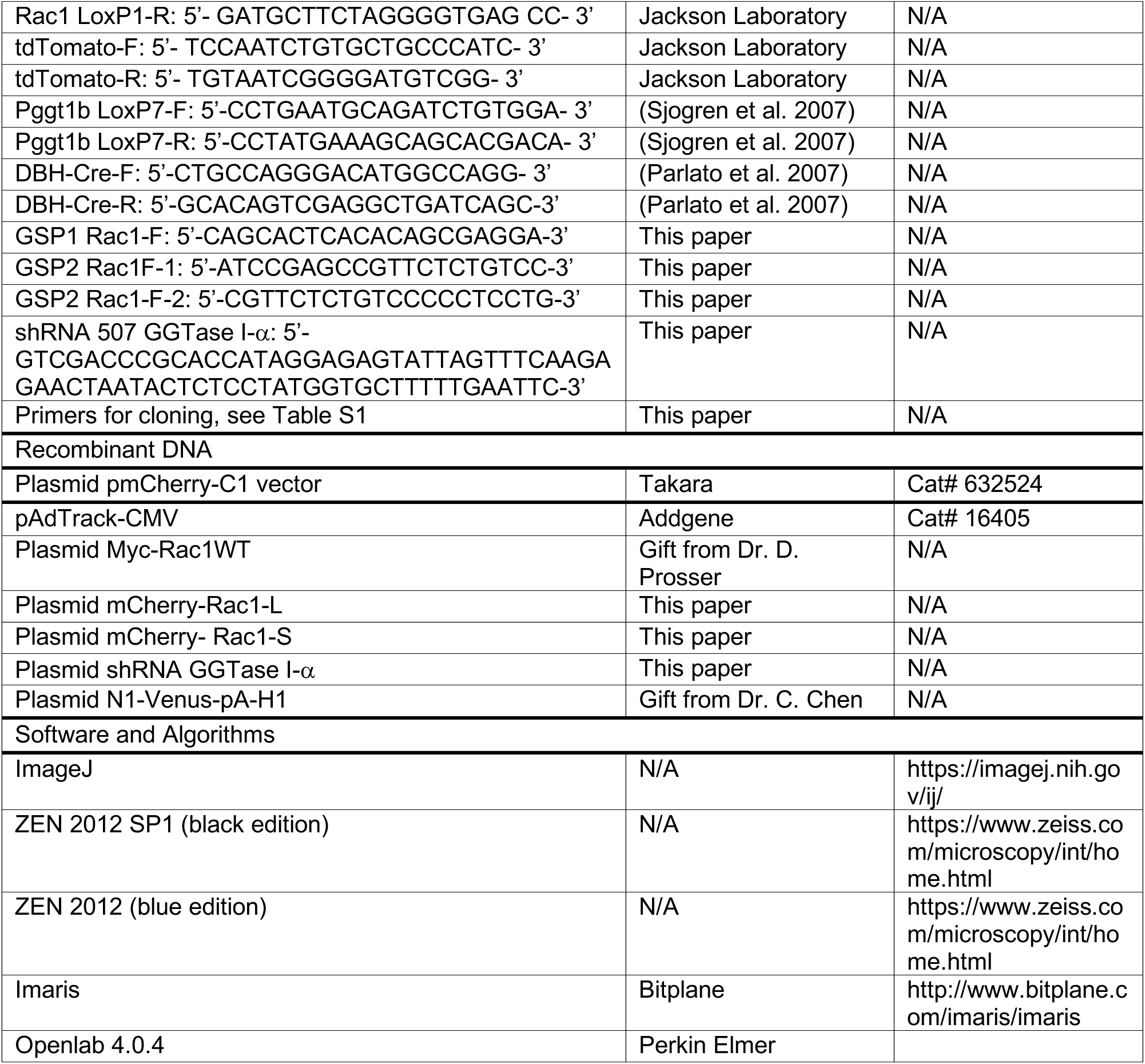

### CONTACT FOR REAGENT AND RESOURCE SHARING

Further information and requests for resources and reagents should be directed to and will be fulfilled by the Lead Contact, Rejji Kuruvilla.

### EXPERIMENTAL MODEL AND SUBJECT DETAILS

#### Materials and Methods

##### Animals

All procedures relating to animal care and treatment conformed to The Johns Hopkins University Animal Care and Use Committee (ACUC) and NIH guidelines. Animals were group housed in a standard 12:12 light-dark cycle. Postnatal day P0.5-P6 pups of both sexes were used for analyses. The following mouse lines were used in this study; *Rac1^fl/fl^* (*Rac1^tm1Djk^/J*) mice (Jackson Laboratory, stock no. 005550), *Rosa-26^tm9(CAG-tdTomato)^* (Jackson Laboratory, stock no. 007909), *Pggt1b^fl/fl^* mice (Sjogren et al., 2007) were generously provided by Dr. Martin Bergö (Karolinska Institutet), *DBH-Cre* mice (Parlato et al., 2007) by Dr. Warren Tourtellotte (Cedars Sinai Medical Center), and *TH-Cre* mice (Gong et al., 2007) by Dr. Charles Gerfen (NIH).

Pregnant Sprague Dawley rats were purchased from Charles River or Taconic Biosciences. Dissociated or explant cultures of sympathetic neurons were established from superior cervical ganglia (SCG) dissected from P0.5 rat pups as previously described (Zareen and Greene, 2009).

##### Neuronal cultures

Sympathetic neurons were harvested from P0.5 Sprague-Dawley rats or *Rac1^fl/fl^* mice and were grown as explant cultures, or dissociated for mass or compartmentalized cultures. Neurons were maintained with high-glucose DMEM media supplemented with 10% fetal bovine serum (FBS), penicillin/streptomycin (1U/ml), and NGF (100 ng/ml). For immunocytochemistry, cells were grown on coverslips coated with poly-d-lysine (1 μg/ml; Sigma-Aldrich) and laminin (10 μg/m; Sigma-Aldrich). NGF deprivation was performed in high-glucose DMEM supplemented with 1% FBS with anti-NGF (1:1000) and the caspase inhibitor, boc-aspartyl(O-methyl)-fluoromethylketone (BAF, 50 μM) for 36 hr. For adenovirus infections, neurons were infected with high-titer viruses for 36-48 hr. For electroporation with mCherry-Rac1-L or mCherry-Rac1-S constructs, sympathetic neurons isolated from P0.5 rats were electroporated using P3 Primary Cell 4D-Nucleofector® X Kit (program number: CA-137) according to the manufacturer’s instructions.

### METHOD DETAILS

#### Adenoviral and plasmid constructs

Human Myc-Rac1 coding sequence was fused using PCR to the *Rac1* short or long 3’UTR from RACE clones acquired from compartmentalized cultures of rat sympathetic neurons. The CaaX motif mutation (C189S) was introduced by PCR-based site-directed mutagenesis. Plasmids were verified by DNA sequencing. *Rac1* constructs were sub-cloned into pAdTrack-CMV using SalI and NotI restriction sites. Recombinant adenoviral constructs for Myc-tagged Rac1-long 3’UTR (Rac1-L) or Rac1-short 3’UTR (Rac1-S) were generated using AdEasy adenoviral system and recombinants were transfected into HEK 293 cells using Lipofectamine 3000. Adenovirus for Myc-Rac1-L-C189S was generated by ViGene Biosciences, Inc. based on recombinant plasmid generated by E.S. High titer virus stocks were obtained using Vivapure® AdenoPACK^TM^.

mCherry-Rac1-L and mCherry-Rac1-S plasmids were generated by subcloning myc-Rac1-L and myc-Rac1-S from pAdTrack-CMV using Kpn1 and EcoRV restriction sites into pmCherry-C1, which had been linearized by KpnI and SmaI.

To generate shRNA for GGTase I-α, a custom oligonucleotide with the sequence 5’-GTC GAC CC<u>G CAC CAT AGG AGA GTA TTA GT</u>T TCA AGA GA<u>A CTA ATA CTC TCC TAT GGT GC</u>T TTT TGA ATT C-3’, directed against the GGTase I CDS, was digested with SalI and EcoRI, and then ligated into N1 Venus pA H1 backbone which had been linearized with XhoI and EcoRI. The shuttle vector was derived from pEGFP-N1 (Clontech 6085-1).

#### RT-PCR and qRT-PCR analyses

Cell body and axonal RNA was isolated from rat sympathetic neurons grown in compartmentalized cultures using Trizol. cDNA was generated using Ambion RETROscript kit and transcripts analyzed by PCR using gene specific primers (see **Table S1** for primer sequences).

For Rac1 deletion, sympathetic neuron cultures established from P0.5 *Rac1^fl/fl^* mice were infected with adenoviruses for Cre or LacZ. mRNA was extracted 48 hr post-infection using RNAqueous Micro Total RNA Isolation Kit. To confirm deletion of *Pggt1b*, superior cervical ganglia (SCG) were isolated from P0.5 *Pggt1b^fl/fl^* or *DBH-Cre;Pggt1b^fl/fl^* mice and mRNA isolated as described above. cDNA was prepared using Superscript IV First Strand Synthesis System. Real-time qPCR analysis was performed using TaqMan probes for *Rac1* or *Pggt1b* in a StepOnePlus^TM^ Real-Time PCR Systems (ThermoFisher). Each sample was analyzed in triplicate reactions. Fold change in transcript levels were calculated using the 2^(-ΔΔCt)^ method, normalizing to 18s rRNA transcript.

#### 3’ RACE analysis

Total RNA was isolated from cell bodies or axons of compartmentalized sympathetic neuron cultures and cDNA generated as described above. cDNA generation was done using a hybrid primer containing an oligo-(dT) to enrich for the 3’-end of mRNA and a unique 35-nucleotide sequence, as previously described (Scotto-Lavino et al., 2006). Rac1 3’UTR was then amplified using a primer against the 35-nucleotide sequence, and a *Rac1* CDS specific primer. Non-specific products were removed through a second amplification step using two different *Rac1* CDS specific primers (See **Table S1: Primer Sequences**). PCR products were separated on an agarose gel, and bands purified using NucleoSpin® Gel Clean-Up. Purified bands were cloned into the TOPO TA Cloning System and sequenced in both orientations.

#### Prenylation assays

##### Visualization of protein prenylation

Visualization of protein prenylation in sympathetic neurons was performed as previously described (Gao and Hannoush, 2014), with a few modifications. Briefly, sympathetic neurons grown in mass cultures or compartmentalized cultures were deprived of NGF for 36 hr, and then treated with propargyl-farnesol (isoprenoid analog, 25 μM) in the presence of NGF (50 ng/ml), NGF + GGTI-2133 (75 nM) or anti-NGF (1:1000) for 4 hr. After treatments, cells were washed with ice cold PBS and fixed at room temperature for 10 minutes in 4% PFA/ in Cytoskeleton Buffer supplemented with Sucrose (CBS; 10mM MES pH 6.1, 138 mM KCl, 3mM MgCl, 2mM EGTA, 0.32 M sucrose). After PBS washes, neurons were permeabilized with 0.1% Triton in PBS for 2 min followed by extensive PBS washes to remove detergent. A click reaction cocktail containing 1 mM CuSO_4_, 250 μM Biotin-Azide, and 1 mM TCEP (Tris(2-carboxyethyl) phosphine hydrochloride)) in 100 mM phosphate buffer pH 7.4 (PB) was added to each coverslip and incubated for 1 hr in the dark. Coverslips were thoroughly washed and blocked in 1%BSA/5% goat serum/0.1%Trition X-100 in PBS for 1 hr and then incubated overnight with anti-β-III-tubulin at 4°C. Cells were washed with PBS and incubated for 1.5 hr with Streptavidin-Alexa-488 and secondary antibody in blocking solution at room temperature. To visualize axonal growth cones, Phalloidin-Alexa-546 labeling (1:50) was done during secondary antibody incubation. Neurons were then mounted in Aqueous Mounting Medium containing 100 μg/ml DAPI. Images were acquired using a Zeiss LSM 700 confocal scanning microscope.

Explant cultures were grown for 4-5 days in culture, and cell bodies surgically removed using a scalpel. Isolated axons were treated with propargyl-farnesol (isoprenoid analog, 25 μM) in the presence of NGF (50 ng/ml), NGF + cycloheximide (CHX, 25 μM) or anti-NGF (1:1000) for 6 hr. Axons were then fixed, permeabilized, and newly prenylated proteins visualized as described above. β-III-tubulin immunostaining was performed to visualize axons. An area corresponding to 1 mm^2^ of explants was imaged using an LSM 700 confocal microscope. Images represent z-projections that to ensure full coverage of explants; all images were taken at the same intensity.

##### Biochemical assay for prenylated Rac1

SCG explant cultures were grown for 4-5 days, after which cell bodies were excised, and isolated axons incubated with isoprenoid analog (25 μM) in the presence of NGF (50 ng/ml), NGF + GGTI-2133 (75 nM) or NGF + CHX (25 μM) or anti-NGF (1:1000) for 6 hr. Axons were lysed in 0.1% CHAPS/150 mM KCl/50 mM HEPES buffer with cOmplete Mini protease inhibitor cocktail (Roche) and sonicated on ice. Lysates were concentrated using a 10 kDa Amicon Ultra Filter. Rac1 was immunoprecipitated using mouse anti-Rac1 (1μg) for 4 hr at 4°C. Protein G-agarose beads (40 μl) were added and sample incubated for 4 hr at 4°C. After washes with lysis buffer, beads were resuspended in PB and click reaction performed at 4°C for 1 hr to conjugate TAMRA-PEG azide to the isoprenoid group, as previously described (Nishimura and Linder, 2013). Immunoprecipitates and supernatants were immunoblotted using TAMRA and Rac1 antibodies, respectively. Rac1 immunoblots were stripped and re-probed for p85 for protein normalization. All immunoblots were visualized with ProSignal Dura ECL Reagent and scanned with a Typhoon 9410 Variable Mode Imager (GE Healthcare).

##### GGTase I-α immunoblotting

Sympathetic neurons grown in compartmentalized cultures with NGF on distal axons for 7 days in vitro were lysed in RIPA buffer (50 mM Tris-HCl, 150 mM NaCl, 1 mM EDTA, 1% NP-40, 0.25% deoxycholate with cOmplete Mini protease inhibitor cocktail (Roche). Soma and axon lysates were immunoblotted with anti-GGTase I-α antibody (1:1000). For isolated sympathetic axons, the cell bodies of 4-5 day old explants were mechanically removed and axons incubated without NGF, NGF (50 ng/mL) or NGF with CHX (25 μM) for 6 hr.

Axon lysates were collected in 200 μl of RIPA buffer and immunoblotted for GGTase I-α. Blots were stripped and re-probed for p85 for normalization. To confirm antibody specificity, PC12 cells were electroporated with shRNA plasmid using Cell Line Optimization 4D-Nucleofector^TM^ X-kit per manufacturer protocol with empty vector as control. Protein lysates were collected 48 hr later, and GGTase I-α expression assessed by immunoblotting with anti-GGTase I-α. Immunoblots were stripped and re-probed for α-tubulin as loading control.

#### GGTase I enzymatic assay

Compartmentalized sympathetic neurons were stimulated with NGF (50 ng/ml) applied to distal axons for 30 min, after 48 hr of NGF deprivation. Cell body and axon lysates prepared in 0.2% octyl-β-D-glucopyranoside, 50 mM Tris-HCl (pH 7.5) were incubated with 50 μM ZnCl_2_, 5 mM MgCl_2_, 20 mM KCl, 10 μM dansyl-GCVLL peptide and 10 μM geranylgeranyl pyrophosphate. GGTase I activity was measured by an increase in fluorescence at 460 nm using a Tecan Infinite 200 plate reader (Pickett et al., 1995). GGTase I activity was normalized to total protein using a Pierce^TM^ BCA Protein Assay Kit. To assess the effect of CHX on GGTase I activity, mass cultures of sympathetic neuron were either deprived of NGF or stimulated with NGF (50 ng/ml) in the presence or absence of CHX (25 μM) for 6 hr. Neuronal lysates were prepared in 0.2% octyl-β-D-glucopyranoside, 50 mM Tris-HCl (pH 7.5) and GGTase I activity determined as described above.

#### Live-cell antibody feeding for Trk trafficking

Live-cell antibody feeding to monitor trafficking of surface Trk receptors was performed as previously described (Yamashita et al., 2017). Briefly, cultured rat sympathetic neurons were infected overnight with a doxycycline-inducible FLAG-TrkB:A adenovirus. Neurons were treated with doxycycline (200 ng/mL, 18 hr) to induce FLAG-TrkB:A expression. Surface FLAG-TrkB:A was labeled by incubating neurons with anti-FLAG antibody (1:500) at 4°C in PBS for 30 minutes, in the absence of BDNF. Some cultures were also co-incubated with GGTI-2133 (75 nM) during surface labeling. Excess antibody was washed off with ice-cold PBS and neurons treated with BDNF (50 ng/mL), propargyl-farnesol (isoprenoid analog, 25 μM), and GGTI-2133 (75 nM) as indicated for 4 hr. Neurons were returned to 4°C and quickly washed multiple times in ice-cold acidic buffer (0.2 M acetic acid, 0.5 M NaCl, pH 3.0) to strip surface-bound FLAG antibody. Neurons were fixed in 4%PFA/PBS, permeabilized with blocking solution (0.1%Triton X-100/5% Normal Goat Serum/PBS) and incubated overnight with mouse anti-β-III-tubulin (1:1000). Internalized FLAG-TrkB:A receptors were visualized by incubation with anti-rabbit-Alexa 546 secondary antibody for anti-FLAG and anti-mouse IgG2b-Alexa-647 secondary antibody for β-III-tubulin in blocking buffer. Following immunostaining, newly prenylated proteins were visualized by conjugation of TAMRA-PEG azide as described above using click chemistry. Axons were visualized using β-III-tubulin immunoreactivity. Images representing 0.8 μm slices were acquired using a Zeiss LSM 700 confocal scanning microscope. The same confocal settings were used to acquire all images taken from a single experiment. Intracellular accumulation of FLAG-TrkB:A receptors in axons was quantified as the number of FLAG-immunopositive punctae per μm. Co-localization between FLAG-Trk and isoprenoid analog-labeled proteins was quantified as the number of FLAG punctae that co-localized with TAMRA signal in axons (from a stretch of axons 75-100 μm from the axon tip) and expressed as a percent of the total FLAG punctae in axonal segments. Results are expressed as means ± SEM and expressed relative to the “no ligand” condition. For all imaging, 15-20 axons were analyzed per condition per experiment.

#### Visualization of mCherry-Rac1-L or mCherry-Rac1-S in sympathetic neurons

Neurons electroporated with mCherry-Rac1-L or mCherry-Rac1-S were grown on poly-D-lysine and laminin coated coverslips for 36 hr and subsequently fixed in 4%PFA/PBS and permeabilized in 0.1%triton/PBS. Neurons were counter-stained with dichlorotriazinylamino fluorescein (DTAF) and mounted in Aqueous Mounting Media containing 100 μg/ml DAPI. Images representing 0.8 μm optical slices were acquired with a Zeiss LSM 700 confocal scanning microscope using automated tiling over several z-stacks to cover whole thickness and length of neuronal cell body and axon. Maximal intensity projections were processed using ImageJ software. Fluorescence intensities for proximal axons were determined by manually tracing 75-100 μm of the axon immediately adjacent to the cell body, and for distal axons by manually tracing 75-100 μm from the axon tip. Average fluorescence intensities were then calculated and normalized to the fluorescent intensities in cell bodies. Total of 19-21 neurons were analyzed per condition per experiment.

#### Neuronal counts

Neuronal counts were performed as previously described (Patel et al., 2015). In brief, torsos of P0.5 mice were fixed in 4%PFA/PBS overnight and cryoprotected in 30% sucrose/PBS for 48 hr. Torsos were then mounted in OCT and serially sectioned (12 μm). Next, every fifth section was stained with solution containing 0.5% cresyl violet (Nissl). Cells in both SCGs with characteristic neuronal morphology and visible nucleoli were counted using ImageJ.

#### Axon growth

For assessing axon growth, neurons isolated from P0.5 rats were grown in Campenot chambers for 7-9 days. Neurons were either completely deprived of NGF or NGF (50 ng/ml) was added only to distal axons. BAF (50 μM) was also included to allow assessment of axon growth without the complications of cell death. For compartmentalized inhibition of prenyltransferases, rat sympathetic neurons in compartmentalized cultures were treated with GGTI-2133 (75 nM) or FTI-277 (100 nM) added either exclusively to cell body or distal axon compartments. Phase contrast images of axons were captured using a Retiga EXi camera in 24-hr intervals for 3 days on a Zeiss Axiovert 200 microscope. Axon growth rate was measured using Openlab 4.0.4 for an average of 30-60 axons per condition. Axons were fixed in 4% PFA and stained with β-III-tubulin for representative images following experiments.

Compartmentalized cultures from P0.5-P6 *Rac1^fl/fl^* mice were infected with adenoviruses expressing GFP, Cre, Cre + Rac1-L, Cre + Rac1-S, or Cre + *Rac1*-L-C189S after axons had extended into the side compartments (7-10 days *in vitro*). Axon growth was then assessed in response to axon-applied NGF (50 ng/ml) in 24 hr intervals for 48 hr, as described above.

#### Immunostaining

P0.5 mouse sections (12 μm) from *TH-Cre;TdTomato* reporter mice were permeabilized with 0.5% Triton X-100 in Phosphate Buffered Saline (PBS) and blocked using 3%BSA/PBS/0.5% Triton X-100. Sections were then incubated with rabbit anti-GGTase I-α antibody (1:200) overnight. Following PBS washes, sections were incubated with anti-rabbit Alexa-488 secondary antibody (1:200). Sections were then washed in PBS and mounted in Fluoromount Aqueous Mounting Medium containing 100 μg/ml DAPI. Images representing 0.8 μm optical slices were acquired using a Zeiss LSM 700 confocal scanning microscope.

P0.5 rat SCGs were grown as explants or dissociated sympathetic neuron cultures on poly-d-lysine and laminin-coated glass coverslips. Neurons were fixed for 30 min. at room temperature in 4% Paraformaldehyde (PFA) in Phosphate Buffered Saline (PBS). Coverslips were washed with PBS and blocked in 3%BSA/0.1%Triton X-100/PBS or 5%Goat Serum/1%BSA/0.1% Trition X-100/PBS for 1 hr. Explants or dissociated neurons were incubated overnight with β-III-tubulin (1:500) or GGTase I-α (1:200). After washes with PBS, neurons were incubated with anti-rabbit-488 or anti-mouse-546 secondary antibodies. In immunostaining for GGTase I-α, 5-(4,6-dichlorotriazinyl) aminofluorescein (DTAF), a reactive dye that labels amines in proteins, was added at 1:10,000 (stock 10mg/ml) for 1 hr as a counterstain, prior to adding the primary antibody. Neurons were then mounted in Aqueous Mounting Medium containing 100 μg/ml DAPI. Images representing 0.8 μm optical slices were acquired using a Zeiss LSM 700 confocal scanning microscope.

#### iDISCO and whole mount immunostaining

iDISCO-based tissue clearing for whole mount immunostaining of organs from P0.5 mice was performed as previously described (Renier et al., 2014). Briefly, kidneys and hearts were fixed in 4%PFA/PBS, then dehydrated by methanol series (20-80%) and incubated overnight in 66% dichloromethane (DCM)/33% methanol. Samples were then bleached with 5% H_2_O_2_ in methanol at 4°C overnight, then re-hydrated and permeabilized first with 0.2%TritonX-100 followed by overnight permeabilization with 0.16% TritonX-100/20%DMSO/0.3M glycine in PBS. Samples were incubated in blocking solution (0.17% TritonX-100/10% DMSO/6% Normal Goat Serum in PBS) for 8 hr, and then incubated with rabbit-anti-TH (1:200) in 0.2% Tween-20/0.001% heparin/5% DMSO/3% Normal Goat Serum in PBS at 37°C for 48 hr. Samples were then washed with 0.2% Tween-20/0.001% heparin in PBS and incubated with anti-rabbit Alexa-546 secondary antibody (1:400) in 0.2% Tween-20/0.001% heparin/3% Normal Goat Serum in PBS. After 48 hr, organs were extensively washed with 0.2% Tween-20/0.001% heparin in PBS and dehydrated in methanol. Samples were cleared by successive washes in 66% DCM/ 33% methanol, 100% DCM and 100% Dibenzyl Ether. Organs were imaged on a lightsheet microscope (LaVision BioTec Ultra Microscope II). Imaris was used for 3D manipulations. Total axon lengths and number of branch points were quantified using Imaris Filament Tracer and normalized to total organ volume.

### QUANTIFICATION AND STATISTICAL ANALYSIS

Sample sizes were similar to those reported in previous publications (Bodmer et al., 2011; Yamashita et al., 2017; Andreassi et al., 2010). Data were collected randomly. For practical reasons, analyses of neuronal cell counts and axon innervation in mouse tissues were done in a semi-blinded manner such that the investigator was aware of the genotypes prior to the experiment, but conducted the staining and data analyses without knowing the genotypes of each sample. All Student’s t tests were performed assuming Gaussian distribution, two-tailed, unpaired, and a confidence interval of 95%. One-way or two-way ANOVA analyses with post hoc Tukey test were performed when more than two groups were compared. Statistical analyses were based on at least 3 independent experiments, and described in the figure legends. All error bars represent the standard error of the mean (s.e.m.).

