## Supplemental Data for "Prenylation of axonally translated proteins controls NGF-dependent axon growth"

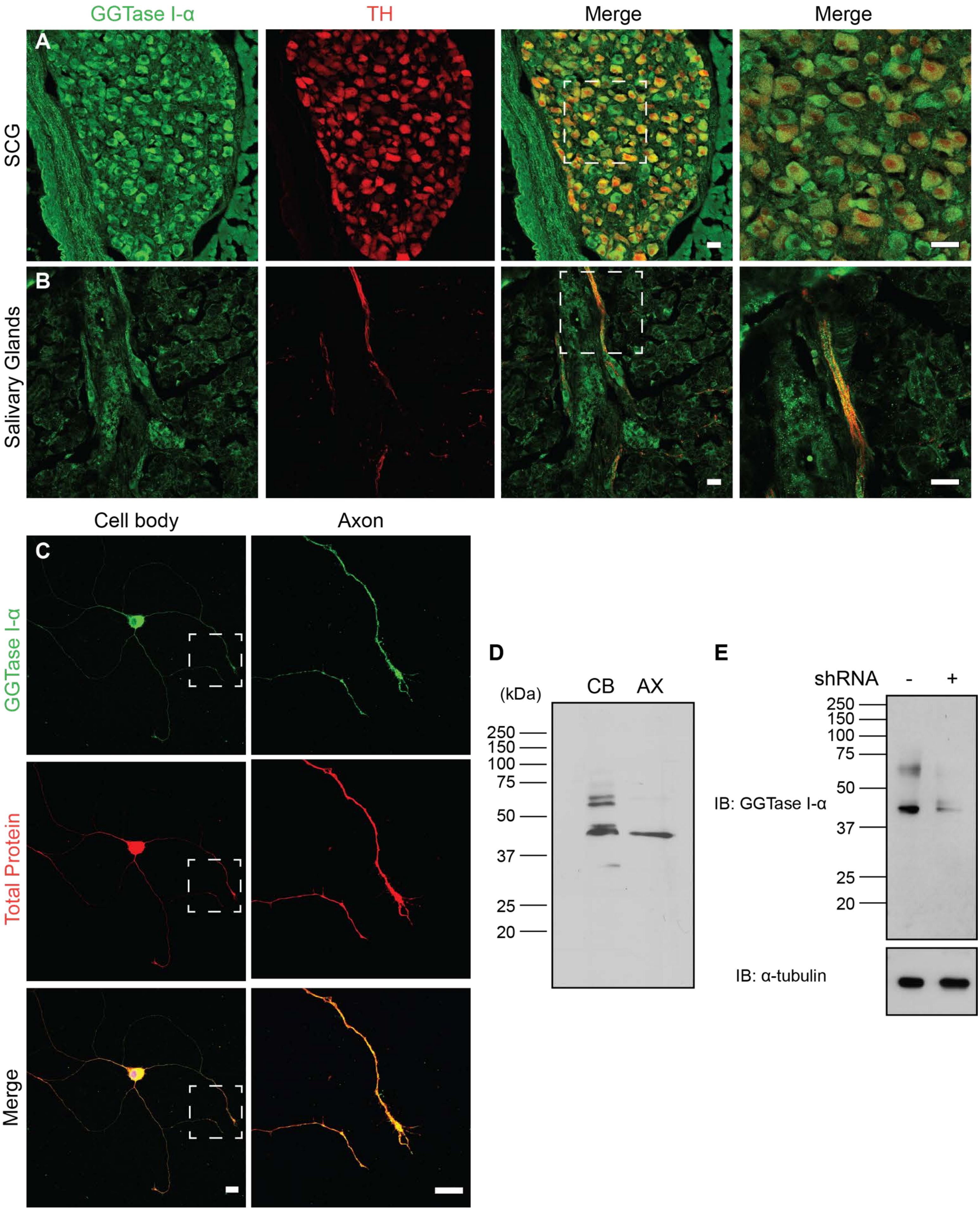
GGTase I-α
TH
Merge
Merge
A
SCG
B
Salivary Glands
Cell body
Axon
C
GGTase I-α
Total Protein
Merge
D
(kDa)
CB
AX
250
150
100
75
50
37
25
20
E
shRNA
-
+
250
150
100
75
50
37
25
20
IB: GGTase I-α
IB: α-tubulin

**Figure S1. Prenyltransferase α-subunit localization in cell bodies and axons of sympathetic neurons**. **Related to Figure 1.** (**A-B**) Immunohistochemistry shows prenyl transferase-α subunit (GGTase I-α) expression in the superior cervical ganglia (SCG) and sympathetic axons innervating the salivary glands in *TH-Cre;TdTomato* reporter mice at postnatal day 5 (P5). Scale bar: 25 μm. (**C**) Prenyl transferase-α localization in sympathetic neuron cell body and axon in culture. Neurons were counter-stained with a total protein stain, DTAF. Right panels are magnifications of the areas in the insets. Scale bar: 10 μm. **(D)** Western blot analysis of prenyl transferase-α expression in cell bodies (CB) and axons (AX) from compartmentalized sympathetic neuron cultures. **(E)** Western blot of prenyl transferase-α and α-tubulin expression in PC12 lysates transfected with shRNA against prenyl transferase-α or empty vector. α-tubulin expression is shown as loading control.

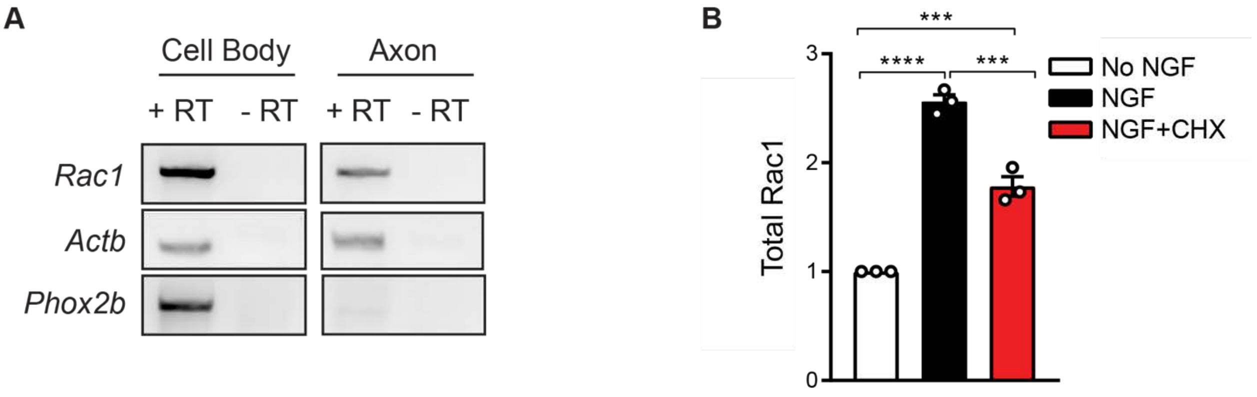


**Figure S2. Axonal synthesis of Rac1. Related to Figure 5. (A)** RT-PCR analysis shows *Rac1* mRNA and *Actb* (β-actin), a known axonal transcript, in axonal lysates from compartmentalized sympathetic neuron cultures. *Phox2b* is a cell body-specific transcript. (**B**) NGF (50 ng/ml) enhances Rac1 protein in isolated sympathetic axons that is reduced by CHX (25 μM) treatment. Quantification of total Rac1 protein in axons normalized to p85. Results are mean ± s.e.m from n=3 experiments, 50-70 explants pooled per condition for each experiment,***p<0.001, ****p<0.0001, one-way ANOVA Tukey's multiple comparisons.

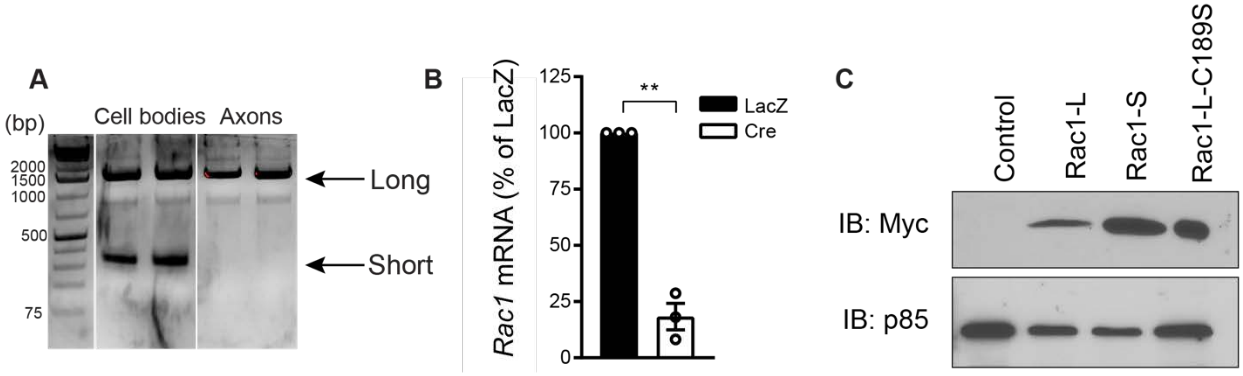


**Figure S3. Sympathetic neurons express two major *Rac1* isoforms. Related to Figure 6.** (**A**) 3' RACE on mRNA isolated from cell bodies or distal axons of sympathetic neurons using a second primer set. **(B)** RT-qPCR of *Rac1* mRNA levels in *Rac1*$^{fl/fl}$ sympathetic neuron cultures infected with adenovirus expressing LacZ or Cre. Values are normalized by levels of *18S* rRNA and expressed as percentage of LacZ-infected neuron values. Data are presented as mean ± s.e.m. from n=3 experiments, **p<0.01, one-sample t-test. (**C**) Western blotting analysis of Myc and p85, as loading control, in sympathetic neuron cultures infected with adenoviruses for Myc-Rac1-long 3'UTR (Rac1-L), Myc-Rac1-short 3'UTR (Rac1-S) or Myc-Rac1-L with a mutated CaaX motif (Rac1-L-C189S).

A

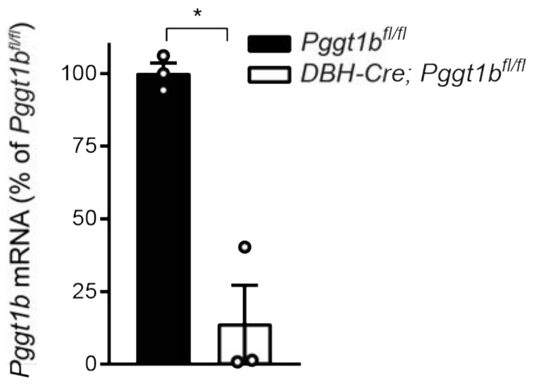


**Figure S4. Deletion of GGTase I during sympathetic neuron development. Related to Figure 7. (A)** SCGs were harvested from P0.5 *DBH-Cre;Pggt1b*$^{fl/fl}$ mice and *Pggt1b*$^{fl/fl}$ littermates. Levels of *Pggt1b* mRNA were analysed by RT-qPCR, normalised by *18S* rRNA and are calculated as percentage of the *Pggt1b*$^{fl/fl}$ values. Data are presented as mean ± s.e.m. from n=3 mice per genotype, *$p<0.05$, one-sample t-test.

**Table S1: Primer Sequences:**

| Application | Primer Name | Primer Sequence |
|---|---|---|
| RT-PCR | Rac1F<br>Rac1R | 5'- CTACCCGCAAACAGACGTG-3'<br>5'- CCATTTTCTGAGCAAAGCGT- 3' |
| RT-PCR | Acbt (β-actin)F<br>Acbt (β-actin)R | 5'- ATGGATGACGATATCGCTGCG– 3'<br>5'- GGTGACAATGCCGTGTTCAAT- 3' |
| RT-PCR | Phox2bF<br>Phox2bR | 5'- AGGCCAGTGGCTTCCAGTAT- 3'<br>5'- TGTCTCAGCGAAGACCCTCT- 3' |
| RT-PCR | Fnta (GGTase I-α)F<br>Fnta (GGTase I-α)R | 5'-AGTTTGGCACCATAGGAGAG-3'<br>5'-ACTCCTGAATGACCCACTGT-3' |
| RT-PCR | Pggt1b (GGTase I-β)F<br>Pggt1b (GGTase I-β)R | 5'-CGAGGTTCTTCATATTTGGG-3'<br>5'-CGCTATGTGTCCACTGTCAT-3' |
| RT-PCR | 18SF<br>18SR | 5'-CGCCGCTAGAGGTGAAATTC-3'<br>5'-TTGGCAAATGCTTTCGCTC-3' |
| Genotyping | Rac1LoxP1F<br>Rac1LoxP1R | 5'-TCC AAT CTG TGC TGC CCA TC-3'<br>5'-GAT GCT TCT AGG GGT GAG CC-3' |
| Genotyping | tdTomatoF<br>tdTomatoR | 5'-ATGGTGAGCAAGGGCGAGG-3'<br>5'-TGTAATCGGGGATGTCGG-3' |
| Genotyping | Pggt1bLoxP7F<br>Pggt1bLoxP7R | 5'-CCTGAATGCAGATCTGTGGA-3'<br>5'-CCTATGAAAGCAGCACGACA-3' |
| Genotyping | DBH-CreF<br>DBH-CreR | 5'-CTGCCAGGGACATGGCCAGG-3'<br>5'-GCACAGTCGAGGCTGATCAGC-3' |
| 3'UTR-Race | GSP1 Rac1F | 5'-CAGCACTCACACAGCGAGGA-3'. |
| 3'UTR-Race | GSP2 Rac1F1 | 5'-ATCCGAGCCGTTCTCTGTCC-3' |
| 3'UTR-Race | GSP2 Rac1F2 | 5'-CGTTCTCTGTCCCCCTCCTG-3' |
| shRNA | shRNA507GGTase I-α | 5'-GTCGACCCGCACCATAGGAGAGTATTA GTTTCAAGAGAACTAATACTCTCC TAT GGTGCTTTTTGAATTC-3' |
| Cloning | Rac1CLLLF<br>Rac13UTR insert NotIR | 5'-GAGAAAATGCCTGCTGTTGTAAATGTCTGAGCCCCT-3'<br>5'-AAGGTTGCGGCCGCGAGGACTCGAGC TCAAGC-3' |
| Cloning | Rac1SLLLF<br>Rac13UTR insert NotIR | 5'-GAGAAAATCACTGCTGTTGTAAATGTCTGAGCCCCT-3'<br>5'-AAGGTTGCGGCCGCGAGGACTCGAGC TCAAGC-3' |
| Cloning | Rac1insertSalIF<br>Rac1commonR | 5'-AGTCCGGTCGACATGGAGCAGAAG-3'<br>5'-AAGGTTGCGGCCGCGAGGACT-3' |
